# Chemo-Mechanical Cues Modulate Nano-Scale Chromatin Organization in Healthy and Diseased Connective Tissue Cells

**DOI:** 10.1101/2021.04.27.441596

**Authors:** Su-Jin Heo, Shreyasi Thakur, Xingyu Chen, Claudia Loebel, Boao Xia, Rowena McBeath, Jason A. Burdick, Vivek B. Shenoy, Robert L. Mauck, Melike Lakadamyali

**Author notes:** co-corresponding authors **Addresses for Correspondence:** *Melike Lakadamyali*, Ph.D., Associate Professor, Department of Physiology, Perelman School of Medicine, University of Pennsylvania, 415 Curie Blvd., Clinical Research Building, Room 764, Philadelphia, PA 19104, *Robert L. Mauck*, Ph.D., Mary Black Ralston Professor of Orthopaedic Surgery and Bioengineering, Department of Orthopaedic Surgery, Perelman School of Medicine, University of Pennsylvania, 36^th^ Street and Hamilton Walk, Philadelphia, PA 19104.

## Abstract

Microscale changes in tissue environment are translated to changes in cell behavior and phenotype, yet the mechanisms behind how these phenotypic changes occur are poorly understood. Here, we describe and model chromatin, which stores genetic information within the cell nucleus, as a dynamic nanomaterial whose configuration is modulated by chemo-mechanical cues in the microenvironment. Our findings indicate that physiologic chemo-mechanical cues can directly regulate chromatin architecture in progenitor cell populations. Via direct experimental observation and modeling that incorporates phase transitions and histone methylation kinetics, we demonstrate that soft environmental cues drive chromatin relocalization to the nuclear boundary and compaction. Conversely, dynamic stiffening attenuates these changes. Interestingly, in diseased human fibrous tissue cells, this link between mechanical inputs and chromatin nano-scale remodeling is abrogated. These data indicate that chromatin dynamics and plasticity may be hallmarks of disease progression and targets for therapeutic intervention.

## Introduction

Tissue resident cells (including tissue-resident progenitors) continuously sense changes in their chemo-physical environment and use this information to maintain their phenotype and tissue homeostasis^1,2^. Injury or degeneration of load bearing fibrous connective tissues (e.g., tendons, ligaments, meniscus) results in changes in tissue structure and mechanics ^3–5^. These changes can be seen at the bulk material level, as well as with the emergence of local material micro-domains with altered mechanics harboring cells of aberrant phenotypes^5^. However, it remains unclear whether the altered chemo-physical environments in these degenerating tissues drive a change in the nano-scale material organization of the genome that promulgates this aberrant state, or whether these changes occur in a cell autonomous fashion, and this in turn alters the local microenvironment. It is thus essential that we elucidate how native cells respond and adapt to their chemo-physical environment in order to understand normal physiology and treat pathology in these connective tissues.

Ultimately, phenotypic changes in response to chemo-physical cues are linked to the nano-scale material state of the chromatin which in turn modulates accessibility and transcriptional activity at specific loci^6,7^. Since the functional consequences of an altered chromatin architecture occur at the nano-scale (on the order of 10-100 nm)^7^, it has been challenging to study this phenomenon using conventional fluorescence microscopy. While high throughput chromosome conformation capture based methods have provided valuable information about genome organization, such methods typically require a large population of cells, obscuring cell to cell heterogeneity^7,8^. Recently, a number of super-resolution microscopy techniques have emerged that can overcome the diffraction limitations of conventional microscopy. Indeed, super-resolution microscopy [e.g., stochastic optical reconstruction microscopy (STORM) with a subdiffraction spatial resolution of ∼20-30 nm] allows one to address the resolution gap between conventional fluorescence and electron microscopy^7,8^. Recently, using this imaging technique, we obtained high-resolution images of nano-scale chromatin structure of stem cells^7^, and demonstrated that this nano-scale chromatin organization is cell-type specific and that nucleosome-enriched regions (termed “clutches”) increase in size and density with stem cell differentiation^7^.

To extend this work, here we took advantage of these recent advances to determine how biophysical cues regulate spatial nano-scale chromatin organization during progenitor cell differentiation and native tissue degeneration. For this, we explored human mesenchymal stem cell (hMSC) and human tenocyte (hTC) chromatin nano-material architecture, and investigated how chromatin nano-structure changed in response to physiologic and pathologic chemo-physical cues (e.g., soft substrates^9^ or hypoxic conditions^10^) using super-resolution microscopy. Our findings show that exogenous chemo-physical perturbations rapidly and reversibly regulate spatial chromatin organization and localization at the nano-scale level in both hMSCs and hTCs through changes in histone modifications. Further, we found that tendon development and degeneration altered tenocyte nano-scale chromatin structure and mechano-responsivity. Collectively, these data support that alterations in the chemo-physical environments of fibrous connective tissues with development or degeneration impact nano-scale chromatin organization and may inform new therapeutic strategies for treating connective tissue pathologies.

## Results

### Substrate Stiffness Alters Nano-scale Chromatin Organization in Human MSCs

Given that substrate stiffness plays an important role in determining stem cell fate^11^, and tissues have a wide range of stiffnesses that change with disease^12,13^, we first assessed how this physiologically and pathologically relevant biophysical cue impacts genome organization at the nano-scale level in human MSCs (hMSCs). To do this, hMSCs were plated on chambered cover glass (glass, ∼70 GPa), 30 kPa (Stiff), or 3 kPa (Soft) methacrylated hyaluronic acid hydrogels (MeHA) modified with RGD to promote cell adhesion^14^ (**Fig. 1a**), followed by 2 days of culture in basal cell growth media. Super-resolution images of histone H2B provided a genome wide view of chromatin organization in hMSC nuclei cultured on these substrates (**Fig. 1b**). These images revealed histone nanodomains, which could not be observed with conventional microscopy^7^. Moreover, there were striking differences in the nuclear localization patterns of these chromatin nanodomains, depending on substrate stiffness.

**Figure 1.**
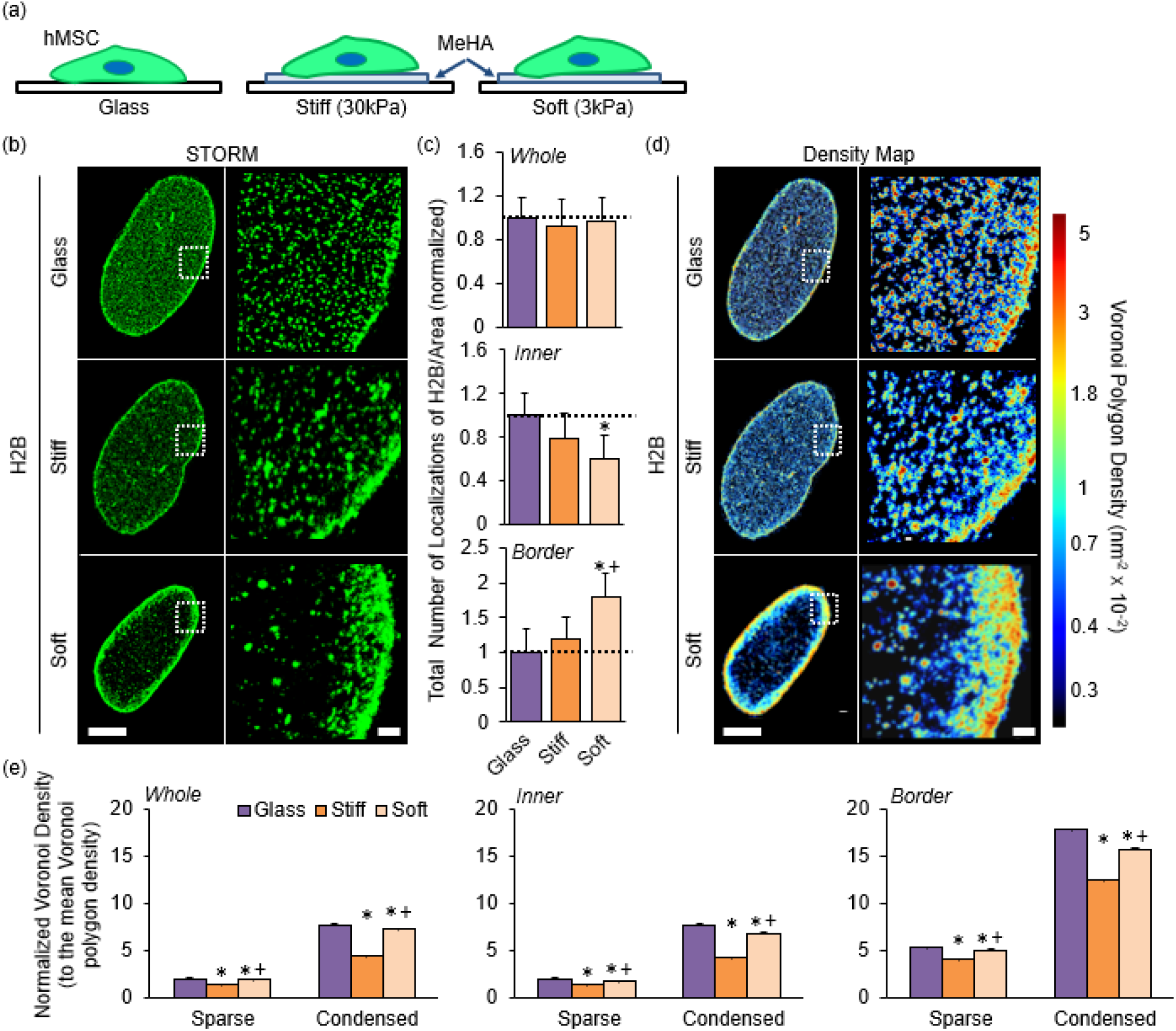
Substrate stiffness regulates histone H2B localizations in human MSCs. (a) Schematic showing hMSCs cultured on cover glass (glass) (∼70 GPa), stiff (30 kPa), or Soft (3 kPa) methacrylated hyaluronic acid (MeHA) hydrogels for 2 days. (b) Representative STORM super-resolution images of H2B in hMSCs cultured on glass, stiff, or soft hydrogels [scale bars = 5 μm (left) and 300 nm (right)]. (c) Quantification of the total number of localizations of H2B per unit area calculated for the whole, the border, or the inner region of the hMSC nuclei on different substrates, showing the enhanced localization of H2B to the nuclear border on soft substrates. The values have been normalized to that on Glass substrate. H2B was labeled with primary and secondary antibodies and STORM super-resolution images of H2Bs were obtained using the photo switchable AlexaFluor647 conjugated to the secondary antibody. Localizations correspond to individual detections of the AlexaFluor647 in the STORM super-resolution images of H2Bs and the total number of localizations is proportional to (but due to the variable labeling stoichiometry and repeated fluorophore blinking, not equal to) the total number of H2B proteins (n = 5 nuclei/group, *: p<0.05 vs. Glass, +: p<0.05 vs. Stiff) (d) STORM super-resolution images of H2B rendered as Voronoi density map based on Voronoi tessellation of H2B localizations showing substrate dependent changes to chromatin spatial localization and compaction. Voronoi polygons were color coded according to their area with red corresponding to small Voronoi polygons (i.e. high H2B density) and blue corresponding to large Voronoi polygons (i.e. low H2B density) according to the color scale bar. (e) Quantification of the Voronoi polygon density (inverse of the Voronoi polygon area) in two compartments: spare euchromatin and condensed heterochromatin calculated for the whole, the border, or the inner region of human MSC nuclei on different substrates. The sparse euchromatin compartment corresponds to 0-40th percentile of the cumulative Voronoi polygon density plot and the condensed heterochromatin compartment corresponds to 41-80th percentile of the cumulative Voronoi polygon density plot averaged over the multiple hMSC nuclei for each substrate. The density of Voronoi polygons in each nucleus has been normalized to the mean Voronoi polygon density in that nucleus to account for any variability in the H2B detections per nucleus (n = 5 nuclei/group, *: p<0.05 vs. Glass, +: p<0.05 vs. Stiff).

To quantify these differences, we used Voronoi tessellation-based image segmentation^7,15^. Voronoi segmentation revealed that while Histone-H2B (H2B) localizations clustered to form discrete and spatially separated nanodomains in hMSC nuclei on all three substrates, these domains were distributed more evenly throughout the nucleus on glass and on stiff substrates compared to soft substrates. (**Fig. 1b**). Surprisingly, H2B nanodomains were mainly sequestered at the nuclear periphery on soft substrates (**Fig. 1b**). Indeed, analysis of localization density revealed that the H2B localizations were enriched at the nuclear periphery (defined as 15 ∼ 20% of the outer area from the nuclear border determined by the image intensity profile across the nucleus) compared to the nuclear interior on soft compared to stiff substrates or glass (**Fig. 1c and sFig 1a-b**). These changes in H2B localization were not due to changes in nuclear volume, as cells grown on different substrates had similar nuclear volume (**sFig 2**). In addition, the total density of H2B localizations throughout the nucleus was similar under the three conditions, suggesting that the total number of visualized H2B molecules was the same (**Fig 1c**). Hence, these results suggest that chromatin as a nanomaterial undergoes spatial reorganization in response to substrate stiffness.

Given the dramatic re-localization of chromatin to the nuclear periphery, which is known to contain silenced, constitutive heterochromatin^16,17^, we next asked if the condensation and compaction state of chromatin was different on the different substrates. To address this question, we used a heatmap representation of the super-resolution images based on Voronoi polygon size (**Fig. 1d**). Since the size of Voronoi polygons is inversely proportional to the localization density, smaller polygons represent denser, more compact chromatin. The H2B heat maps indeed showed visual differences in how the condensed chromatin (red in **Fig. 1c**) was distributed throughout the nucleus on the different substrates. In particular, dense chromatin was somewhat enriched at the nuclear periphery, but also localized throughout the hMSC nucleus on glass substrates (**Fig. 1d**). Conversely, dense chromatin was highly enriched at the nuclear periphery on soft substrates. By placing a threshold on Voronoi polygon density (inverse of Voronoi polygon size) (**sFig. 1c**), we split chromatin into two compartments, <u>sparse</u> euchromatin having low Voronoi density and <u>condensed</u> heterochromatin having high Voronoi density (**sFig. 1c**). This quantification revealed that, compared to glass, both chromatin compartments were less compact on stiff substrates independent of whether the chromatin at the nuclear periphery or interior was analyzed. Interestingly, on soft substrates, both chromatin compartments underwent re-compaction to a similar level as the glass substrate (**Fig. 1c**). Taken together, these data suggest that substrate stiffness regulates nano-scale chromatin spatial organization and compaction in hMSCs.

### Substrate Stiffness Regulates Histone Methylation Status in Human MSC Nuclei

Histone post-translational modifications play crucial roles in regulating gene activation and repression^18–20^, and different modifications correlate with open or closed chromatin. In particular, H3 lysine 4 trimethylation (H3K4me3) is a marker of active euchromatin whereas histone H3 lysine 27 trimethylation (H3K27me3) is associated with heterochromatin and gene repression^21,22^. Having established that substrate stiffness alters chromatin organization and condensation in hMSCs, we next asked if substrate stiffness also leads to changes in the localization and condensation state of active chromatin marked with H3K4me3 and repressed chromatin marked with H3K27me3. We thus labeled hMSCs with antibodies against these two specific histone modifications and obtained super-resolution images under the three substrate conditions. Heat maps of H3K4me3 and quantitative analysis showed that, compared to hMSCs cultured on glass substrates, H3K4me3 levels were slightly increased in hMSCs cultured on stiff substrates, while these levels were significantly lower in hMSCs cultured on the soft substrates (**Fig. 2a,b**). These data are consistent with the H2B condensation analysis above and suggest that stiff substrates may lead to de-condensation regulated by increases in H3K4me3 in hMSCs (**Fig. 1b-e**). Conversely, the amount of H3K27me3 was lower on stiff substrates (in particular at the nuclear interior) and was enhanced on soft substrates (particularly at the nuclear periphery) compared to glass substrates (**Fig. 2c,d**). These results indicate that increased peripheral chromatin localization and increased chromatin condensation on soft substrates is correlated with increased levels of H3K27me3, suggesting that soft substrates may lead to gene suppression (**Fig. 1b-e**).

**Figure 2.**
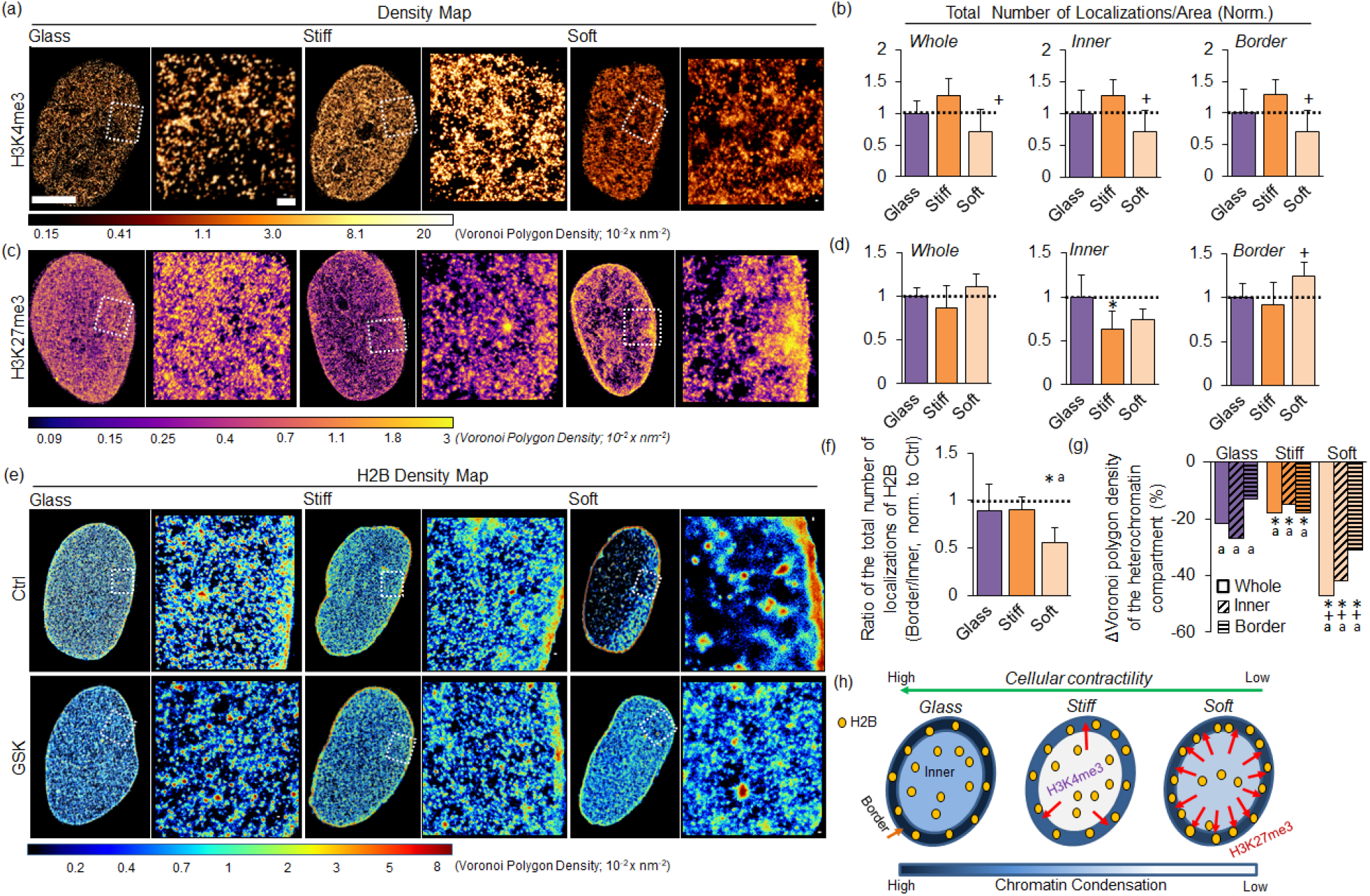
Substrate stiffness-dependent histone methylation in human MSCs. Representative STORM super-resolution images of (a) H3K4me3 or (c) H3K27me3 rendered as a density map based on Voronoi polygon density, with yellow corresponding to high Voronoi polygon density and black corresponding to low Voronoi polygon density according to the color scale bar showing the nanoscale spatial distribution of histone methylation in hMSCs on different substrates (glass, stiff, or soft MeHA hydrogels) (scale bars = 5 μm (left) and 300 nm (right)). Quantification of the total number of (b) H3K4me3 or (d) H3K27me3 localizations per unit area on the whole, the border, or the inner region of the hMSC nucleus on different substrates. The values were normalized to that on Glass substrate (n = 5 nuclei/group, *: p<0.05 vs. Glass, +: p<0.05 vs. Stiff). (e) Representative STORM super-resolution images of H2B rendered as a density map based on Voronoi polygon density with red corresponding to high Voronoi polygon density and blue corresponding to low Voronoi polygon density according to the color scale bar in hMSC nuclei on different substrates (glass, stiff, or soft MeHA hydrogels) with or without GSK343 (a selective inhibitor of EZH2) treatment (Ctrl/GSK). (f) Changes to H2B localization between the nuclear border and inner region of nucleus with and without GSK treatment quantified as the ratio of the number of localizations of H2B per unit area at the nuclear border to the total number of localizations of H2B per unit area at the inner region of the nucleus. The values shown are for GSK treatment, which have been further normalized to the Ctrl group (n = 5 nuclei/group, *: p<0.05 vs. Glass, +: p<0.05 vs. Stiff). (g) Changes to the condensation level of the heterochromatin compartment in GSK treated hMSCs compared to Ctrl hMSCs. The Voronoi polygon density of the condensed heterochromatin compartment in GSK treated hMSCs is shown normalized to control hMSCs, showing a decrease in chromatin condensation of heterochromatin with GSK treatment (n = 5 nuclei/group, a: p<0.05 vs. Ctrl, *: p<0.05 vs. Glass, +: p<0.05 vs. Stiff). (h) Schematics showing substrate stiffness regulates nanoscale spatial chromatin organization and condensation mediated by histone methylation in hMSCs (Yellow circles: H2Bs, Red allows: H2B re-localization to the nuclear border).

Having observed that substrate stiffness regulates chromatin organization at the nano-scale and that substrate dependent chromatin remodeling is correlated to changes in the levels of histone methylation marks, we next investigated the role of the methyltransferase EZH2 (Enhancer of zeste homolog 2, which catalyzes H3K27me3^23,24^) in substrate stiffness mediated chromatin re-organization in hMSCs. To this end, we used GSK343, a selective inhibitor of EZH2^25,26^ to treat hMSCs during the culture period. H2B STORM images, heatmaps and quantitative Voronoi analysis revealed that EZH2 inhibition did not substantially alter H2B spatial organization under stiff or glass substrates, but did lead to a slight de-condensation of both sparse euchromatin and condensed heterochromatin compartments (**Glass and Stiff, Fig. 2e, g, sFig.4-5**). On the other hand, on soft substrates (Soft), EZH2 inhibition reversed the previously observed phenotype of chromatin sequestration at the nuclear periphery (**Soft Fig. 2e, f)**, and led to a substantial de-condensation of both sparse and condensed chromatin compartments (**Fig. 2e, g, sFig.3-4**). Conversely, when Lysophosphatidic Acid (LPA, a G-protein agonist that increases cellular contractility^27,28^) was applied, we observed increases in the condensation state of both sparse and condensed chromatin compartments on all substrates, with the most prominent changes observed in hMSC nuclei on soft substrates (**sFig.5 and 6**). These data suggest that altered physical forces and cellular contractility on soft substrates may play important roles in chromatin translocation to the nuclear periphery in hMSCs. Overall, our results show that substrate stiffness regulates nano-scale chromatin spatial organization and chromatin condensation through changes in cellular contractility and alterations of histone methylation status in hMSCs (**Fig.2h**).

### Theoretical model predicts the formation of nuclear border heterochromatin domains with increased methylation

The formation of heterochromatin has previously been proposed to be a result of liquid-liquid phase separation of chromatin^29,30^. As noted above, specific epigenetic marks are known to correlate with condensed heterochromatin or open euchromatin. Yet, the effects of epigenetic regulators on chromatin organization are not fully understood and whether kinetics of acetylation and methylation can alter chromatin condensation is not known. To explore these points and determine whether the observed impact of substrate stiffness on chromatin condensation is the result of the changes in histone methylation status, we developed a phase-field model that simulates heterochromatin formation [**Fig. 3a** and **Supplementary Model Development and Supplemental Model (SM) Figures 1-3**]. This model incorporates the kinetics of methylation and acetylation (mediated by epigenetic regulators in a substrate-stiffness dependent manner) and interactions of chromatin with nuclear lamina to predict the spatial distribution and volume fraction of heterochromatin.

**Figure 3.**
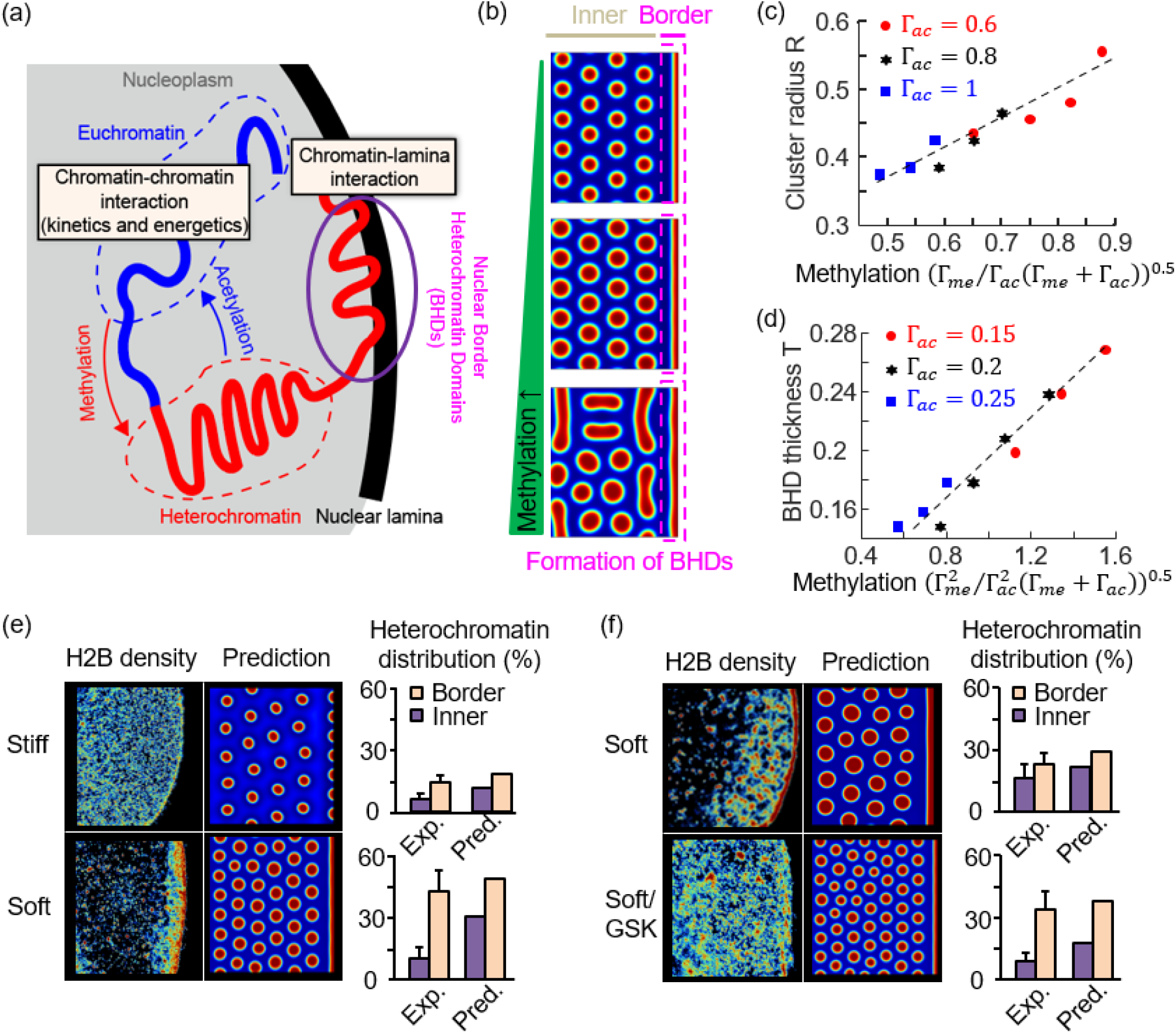
A phase-field model of heterochromatin formation incorporating kinetics of acetylation and methylation recapitulates experimental results. (a) Schematic showing the key features of the phase-field model which captures both the kinetics of acetylation and methylation reactions, and the energetics including chromatin-chromatin and chromatin-lamina interactions. (b) Impact of increasing methylation on the organization of chromatin. Blue and red colors represent euchromatin-rich and heterochromatin-rich phases, respectively. (c) Scaling relation between the size of heterochromatin cluster with the level of methylation. (d) Scaling relation between the thickness of nuclear border heterochromatin domains (BHDs) with the level of methylation. (e) Experimental measurements and model predictions of heterochromatin percentage at the nuclear border and interior for cells cultured on either soft or stiff substrates. (f) Experimental measurements and model predictions of heterochromatin percentage at the nuclear border and interior for cells treated with or without GSK cultured on soft substrates.

The key features in our model include 1) the energetics of chromatin-chromatin and chromatin-lamina interactions specified in Eq. 1 (**Methods**) and 2) the kinetics of chromatin diffusion and chemical reactions described in Eq. 3-4 (**Methods, Fig. 3a-b**). Our model indicates that these two inputs impact the organization of chromatin in different ways. The energetic contributions alone drive the change in particle distribution of heterochromatin into a single large cluster in order to minimize interfacial energy (known as Ostwald ripening^31^). On the other hand, the methylation and acetylation reactions result in interconversion of the eu- and hetero-chromatin phases, resulting in the formation of more interfaces. Faster reactions (higher rate of methylation, *Γ*_*me*_ or acetylation, *Γ*_*ac*_) would thus result in a larger interface area, which competes with the coarsening behavior that minimizes the interfacial energy. Thus, our simulations reveal that the methylation and acetylation reactions suppress the Ostwald ripening, leading to an optimal size of heterochromatin domains (**Fig. 3b**).

This theoretical model agrees with our experimental observations of heterochromatin-rich nanodomains that are stable against coarsening. Further, through our simulations, we found that with increasing levels of methylation, the phase-separated heterochromatin clusters transform from round to lamellar morphologies (**Fig. 3b**) and the size of the heterochromatin clusters increase with the level of methylation (**Fig. 3c**, refer to SI sec.1 for derivation of the scaling relation). In agreement with this prediction, we observed larger heterochromatin clusters when cells were cultured on soft substrates, where we also identified increased levels of methylation (**Fig. 3e**). Finally, the enzymatic reactions also influence the balance between the chromatin-chromatin and chromatin-lamina interactions. Our simulation predicted that more heterochromatin localizes to the nuclear lamina as the level of methylation increases (**Fig. 3b**), consistent with our experimental observations on soft substrates (**Fig. 3e**). The thickness of heterochromatin at the nuclear border also scaled with the methylation levels (**Fig. 3d**, refer to SI sec.2 for derivation of the scaling relation). Our model was also able to accurately predict the decreased localization of chromatin at the nuclear border upon GSK inhibition of EZH2 (**Fig. 3f**), which leads to decreased levels of methylation.

Overall, our model recapitulates the experimental results on how substrate dependent changes in histone methylation levels impact the spatial organization of chromatin within the nucleus and provides support that levels of acetylation and methylation compete with phase separation and have a direct bearing on chromatin organization.

### Rapid Changes in Chromatin Organization in Response to Dynamic Mechanical Cues

The results above indicated that substrate stiffness alters chromatin organization and localization via impacts on cell contractility and downstream methyltransferase activity, but such static, time-invariant mechanical cues do not replicate the dynamic and time varying characteristics of remodeling and actively loaded cellular environments. Hence, we next investigated how dynamic alterations to the biophysical environment impact nano-scale chromatin organization in hMSC nuclei. We used an *in situ* substrate stiffening^32,33^ system to evaluate changes in H2B nano-scale spatial organization and chromatin condensation in hMSCs. Cells were cultured on a stiffening hydrogel system that provided a rapid change in substrate stiffness from a soft (∼3kPa) to a stiff (∼30kPa) mechanical state (**Fig. 4a**)^32^. Consistent with our previous results^32^, hMSCs responded to stiffening immediately with increases in cell area (**sFig. 8a**). As observed in the static soft substrates (3kPa), heat maps of H2B density revealed that chromatin was primarily localized to the nuclear periphery for the first 6 hours after stiffening (**Fig. 4b,c**), followed by a continual redistribution from the nuclear periphery to the nuclear interior between 6→48 hours after stiffening (**Fig. 4b,c**). These changes in chromatin spatial organization were accompanied by a continual decrease in the condensation state of both sparse euchromatin and condensed heterochromatin compartments (**Fig. 4d and sFig. 7b**). Interestingly, changes in condensation started at 2 hours after stiffening, and hence preceded the changes in spatial organization, suggesting that the de-condensation of chromatin likely leads to its spatial remodeling within the nucleus. The amount of decondensation reached a plateau at around 60% of the starting value by 18 hours after stiffening (**Fig. 4d and sFig. 7b**) and lasted up to day 7 (**sFig. 8a-b)**.

**Figure 4.**
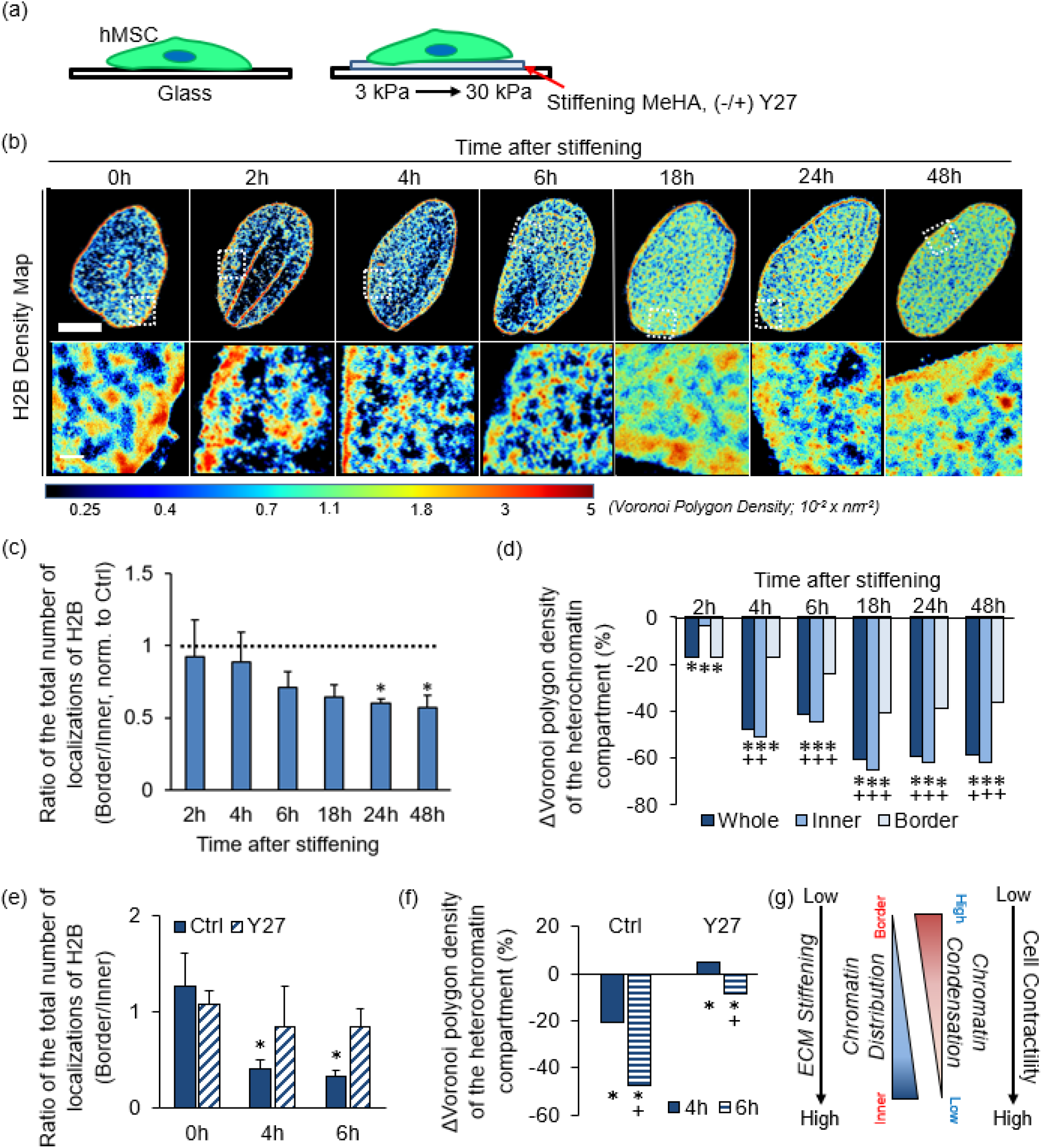
Effect of in situ stiffening on nanoscale chromatin organization in human MSCs. (a) Schematic showing hMSCs cultured on an in situ stiffening hydrogel system providing a rapid change in substrate stiffness from a soft (∼3kPa) to a stiff (∼30kPa) mechanical state with/without addition of the Rock inhibitor Y27632 [(+/−) Y27]. (b) Representative STORM super-resolution images of H2B rendered as a Voronoi polygon density showing a continual redistribution of H2B from the nuclear periphery to the nuclear interior between 0-48 hours after stiffening [bars = 5um (top), 300 nm (bottom)]. (c) Quantification of the ratio of the total number of localizations of H2B at the nuclear border to the total number of localizations of H2B in the inner region of the nucleus. The values have been normalized to the starting value at 0h (n = 5 nuclei/group, *: p<0.05 vs. 0h, +: p<0.05 vs. 2h). (d) Changes in chromatin condensation in condensed heterochromatin compartment in hMSCs between 0-48 hours after stiffening The Voronoi polygon density of the condensed heterochromatin compartment in cells grown on the stiffening hydrogel is shown normalized to the starting Voronoi polygon density at 0h, showing a decrease in chromatin condensation of heterochromatin with time after stiffening (n = 5 nuclei/group, *: p<0.05 vs. 0h, +: p<0.05 vs. 2h). (e) Quantification of the ratio of the total number of localizations of H2B per unit area at the nuclear border to the total number of localizations of H2B at the inner part of the nucleus for hMSCs grown on stiffening hydrogel with/without the Y27 treatment at different times after stiffening, showing that Y27 treatment prevents the relocalization of chromatin from the nuclear border to the nuclear interior upon stiffening (Ctrl/Y27, n = 5 nuclei/group, *: p<0.05 vs. 0h, +: p<0.05 vs. 4h). (f) Changes in chromatin condensation in condensed heterochromatin compartment in hMSCs grown on the stiffening hydrogel with/without Y27 treatment (Ctrl/Y27). The Voronoi polygon density of the condensed heterochromatin compartment in cells grown on the stiffening hydrogel with or without Y27 treatment is shown normalized to the starting Voronoi polygon density at 0h, showing that Y27 treatment prevents the decrease in chromatin condensation of heterochromatin with time after stiffening. (g) Schematic showing how ECM stiffening regulates chromatin distribution and condensation in human MSCs through actomyosin based cellular contractility. ECM stiffening re-distributes chromatin from the border to the inner region of the hMSC nucleus resulting in chromatin decondensation, which requires cellular contractility.

Since our previous study showed that substrate stiffening increased actomyosin-driven contractility of hMSCs within ∼ 4 hours^32^, we next examined the roles of actomyosin-driven contractility in chromatin organization and condensation in response to substrate stiffening. We obtained super-resolution images of H2B in hMSCs cultured on the stiffening MeHA system with or without pretreatment with Y27632 (a ROCK inhibitor, Y27). Consistent with our findings, control hMSCs (without Y27 treatment) altered their spatial organization of chromatin and underwent chromatin de-condensation within 6 hours, while the Y27 treatment blocked this substrate stiffening-mediated chromatin remodeling (**Fig. 4e,f and sFig. 9a**,**b**). These data indicate that ECM stiffening mediated chromatin decondensation and re-distribution from the border to the inner of the hMSC nucleus requires actomyosin based cellular contractility (**Fig. 4g**).

Changes in microenvironmental stiffness represent just one in vivo physical cue that may regulate cell function. To determine if other dynamic physical cues also regulate chromatin nano-organization, we also investigated the effects of fluid-flow induced shear stress (FSS). For this, a custom-PDMS microfluidic chamber was developed to impose FSS to hMSCs seeded on cover glass substrates with varying magnitude (1 ∼ 5 dyne/cm^2^) and duration (0.5 ∼ 2 hours) (**sFig. 10a**). FSS resulted in changes in chromatin distribution with increases in the total number of H2B localizations located at the nuclear periphery and changes in chromatin condensation throughout the nucleus (**sFig. 10b**,**c**). Heat maps of H2B density and quantitative analysis showed that FSS applied at 1 dyne/cm^2^ for 30 mins (1D/0.5h) or for 2h resulted in marked chromatin condensation (**sFig. 10b and d**,**e**). Conversely, higher magnitude FSS [5 dyne/cm^2^] applied for short duration (5D/0.5h)] decreased chromatin condensation. (**Ctrl, sFig. 10b and d**,**e**). Taken together, these data suggest that dynamic biophysical perturbations regulate nano-scale chromatin organizations in a complex and rapid manner in hMSC nuclei.

### Tissue Degeneration and ‘Degenerative’ Chemo-physical Cues Alter Nano-scale Chromatin Organization in Human Tenocytes

Tissue aging and/or degeneration is known to alter the mechanical environment of dense connective tissues, changing both biochemical and biophysical inputs to resident cells, impacting their phenotype^3–5^. Given our results on the impact of biophysical cues on the nano-scale chromatin organization *in vitro*, we next asked how tissue aging and/or degeneration may alter chromatin organization at the nano-scale in native tissue cells. We isolated tenocytes from human tendons from donors that were young and healthy, young and diagnosed with tendinosis (which is a common tendon pathology, associated with repetitive use and mechanical overload^10,34^), or aged and healthy. These human tenocytes were cultured on glass substrates for 48 hours before fixation and super-resolution imaging. Super-resolution heatmaps showed that chromatin was distributed throughout the nucleus in young healthy tenocyte nuclei (Young) and clustered to form distinct and spatially separated H2B nanodomains (**Fig. 5a**). However, and quite strikingly, H2B was more condensed and primarily localized to the nuclear periphery in young degenerative tenocyte nuclei (**Tendinosis, Fig. 5a-b, and sFig. 11a-c**), similar to what we observed for hMSCs cultured on soft substrates. The chromatin organization of degenerative tenocytes was distinct from that of aged tenocyte nuclei. The latter contained less condensed H2Bs throughout the nucleus, similar to hMSCs grown on stiff substrates (**Aged, Fig. 5a-b, and sFig. 11a-c**). These data indicate that both tissue degeneration and aging impact chromatin organization in spatially distinct patterns in human tenocytes.

**Figure 5.**
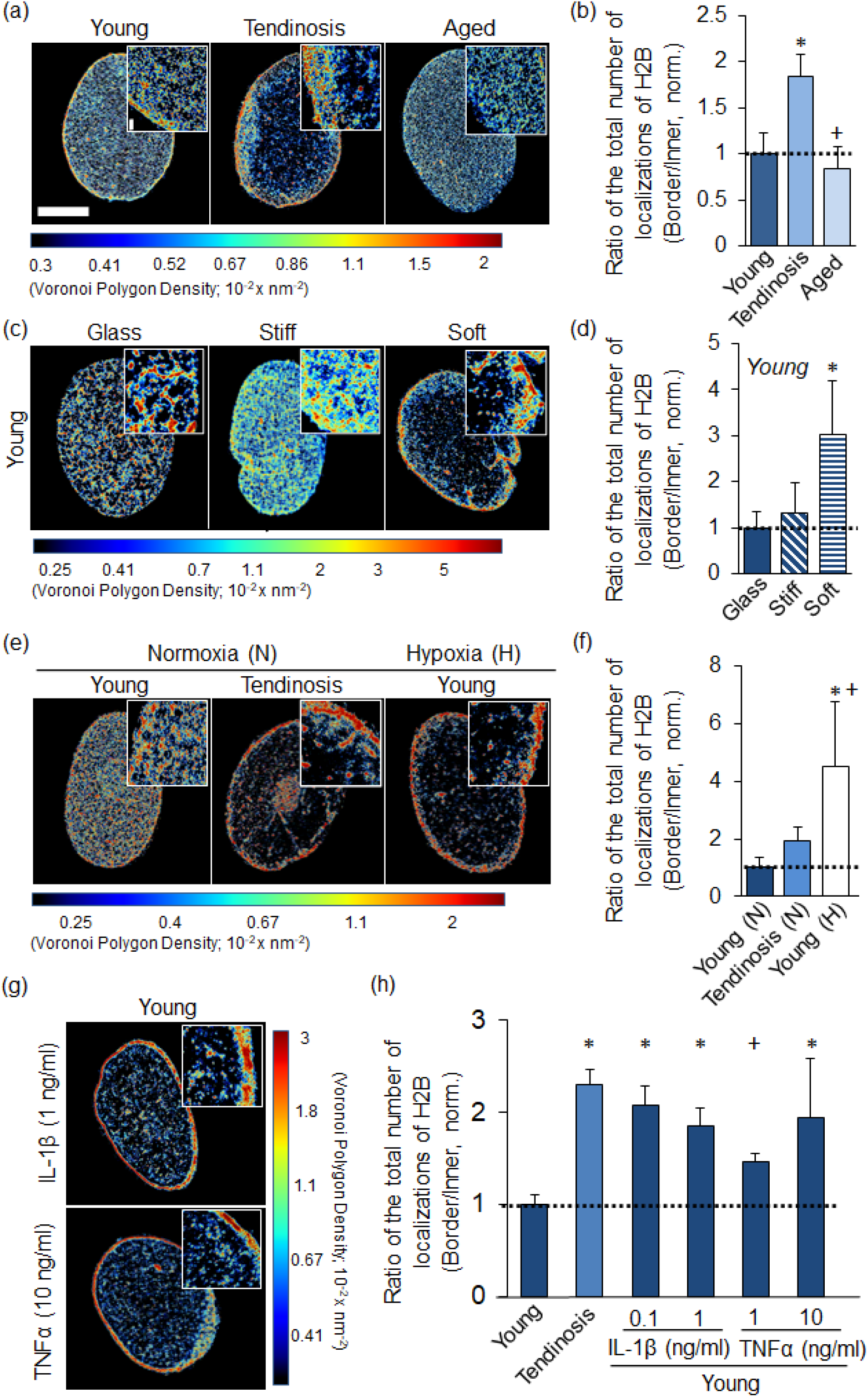
Altered H2B localization in human tenocytes with degeneration or with presentation of ‘degenerative’ chemo-physical cues. (a) Representative STORM super-resolution images of H2B rendered as a density map showing the differences in H2B localization in young and healthy (Young), young and diagnosed with tendinosis (Tendinosis), or aged human tenocytes (Aged). (b) Quantification of the ratio of the total number of H2B localizations per unit area at the nuclear border to the total number of H2B localizations in the inner region of the nucleus. The values have been normalized to young and healthy cells. (n = 5 nuclei/group, *: p<0.05 vs. Young, +: p<0.05 vs. Tendinosis). (c) Representative STORM super-resolution images of H2B rendered as a density map showing changes to H2B localization in human tenocytes cultured on different substrates. (d) Quantification of the ratio of the total number of H2B localizations per unit area at the nuclear border to the total number of H2B localizations in the inner region of the nucleus, showing relocalization of H2B to the nuclear border in human tenocytes grown on soft substrates. The values have been normalized to Glass substrate. (n = 5 nuclei/group, *: p<0.05 vs. glass, +: p<0.05 vs. soft). (e) Representative STORM super-resolution image of H2B rendered as a Voronoi polygon density map showing changes to H2B localization in human young or tendinosis tenocytes under normoxic (N) conditions, or human young tenocytes under hypoxic (H) conditions. (f) Quantification of the ratio of the total number of H2B localizations per unit area at the nuclear border to the total number of H2B localizations in the inner region of the nucleus, showing relocalization of H2B to the nuclear border in young tenocytes under hypoxic conditions. The values have been normalized to young tenocytes grown under normoxic conditions. (n = 5 nuclei/group, *: p<0.05 vs. young healthy tenocyte under the normoxic conditions, +: p<0.05 vs. tendinosis tenocyte under the normoxic conditions). (g) Representative STORM super-resolution images of H2B rendered as a Voronoi polygon density showing changes to H2B localization in human tenocytes cultured on glass substrates with/without (−/+) exposure to IL-1β or TNFα. (h) Quantification of the ratio of the total number of H2B localizations per unit area at the nuclear border to the total number of H2B localizations in the inner region of the nucleus. The values have been normalized to the young human tenocyte control without exposure to IL-1β or TNFα, showing relocalization of H2B to the nuclear border upon treatment with inflammatory cytokines (n = 5 nuclei/group, *: p<0.05 vs. young human tenocyte control, +: p<0.05 vs. tendinosis young human tenocyte). [bars = 5 μm, 300 nm (inset)].

Since we observed different nano-scale chromatin organization in human tenocytes with tissue degeneration or aging, we hypothesized that cues from the degenerative chemo-physical environment (e.g., soft ECM^9^, hypoxia^10^, and inflammation^35^) might result in nano-scale chromatin remodeling, and cause young healthy tenocytes to take on features that are comparable to that seen in aged (Aged) or degenerative (tendinosis) tenocytes. To test this hypothesis, we first cultured human tenocytes isolated from young healthy donors (Young) on glass, stiff (30kPa), or soft (3kPa) substrates for 2 days in basal growth media^4,9^. On glass substrates (Glass), super-resolution heatmaps revealed once again H2B localizations that were clustered to form distinct H2B nanodomains in young healthy tenocyte nuclei (**Fig. 5c**). Interestingly, when these young tenocytes were cultured on stiff substrates (Stiff), chromatin condensation for both sparse and condensed chromatin compartments decreased throughout the nucleus, reaching levels similar to that seen in the aged tenocytes cultured on glass (**Fig. 5c-d, sFig. 12a, b**). Strikingly, chromatin was primarily relocalized to the nuclear periphery on soft substrates, comparable to that seen in degenerative tenocytes (Tendinosis). Chromatin also became more condensed, in particular in the nuclear interior (**Fig. 5c-d, sFig. 12a, b**). Interestingly, the chromatin condensation at the nuclear border decreased for tenocytes grown on soft substrates (**sFig. 12a, b**), unlike the increased condensation seen for degenerative tenocytes (**sFig. 11b, c**), suggesting that disruption of the mechanical microenvironment alone is not sufficient to explain the chromatin remodeling seen in degenerative tenocytes and additional factors are at play. Overall, these data suggest that altered mechanical properties of tissues with tendon aging or micro-damage may drive some of the aberrant changes to chromatin nano-scale organization in disease.

Next, we investigated how changes in oxygen tension impact chromatin organization in young healthy tenocytes, given that previous studies have shown that oxygen levels are altered in degenerative tissues and that hypoxia is a critical regulator of early human tendinopathy^10,36,37^. We seeded young healthy tenocytes on glass substrates for 1 day, followed by 4 days of culture under normoxic (21% O_2_) or hypoxic (1% O_2_) conditions (**Fig. 5e**). Degenerative tenocytes grown under normoxia (Normoxia-Tendinosis) maintained an altered chromatin state consisting of increased chromatin condensation and localization at the nuclear periphery compared to healthy tenocytes grown under normoxia (Normoxia-Young, **Fig. 5f, sFig. 12c, d**). Interestingly, when healthy tenocytes were grown under the hypoxic culture conditions, their chromatin relocalized to the nuclear periphery and underwent increased condensation, similar to that seen in tendinosis tenocytes (Hypoxia-Young, **Fig. 5f, sFig. 12c, d**). These results suggest that, similar to altered mechanical cues, the altered oxygen tension present in tendon tissues as a consequence of injury or degeneration leads to aberrant tenocyte chromatin remodeling.

Because injury and aging are also associated with increased levels of inflammation, we lastly investigated how inflammatory cytokines [e.g., interleukin 1beta (IL-1β), and tumor necrosis factor-alpha (TNFα)] impact nano-scale chromatin organization of young healthy tenocytes. These proinflammatory cytokines are secreted from immune cells that promote tendon inflammation processes in early tendon repair^38,39^. We cultured young tenocytes on glass substrates for 1 day followed by additional culture for 24 hours with or without exposure to IL-1β (0.1 or 1 ng/ml) or TNFα (1 or 10 ng/ml). As observed in tendinopathic nuclei, treatment with proinflammatory cytokines resulted in chromatin reorganization to the nuclear periphery and increased chromatin condensation in young healthy tenocytes (**Fig. 5g, h and sFig. 12e, f**). In the case of TNFα, the magnitude of the observed changes in chromatin organization and condensation was dose dependent and treatment with higher concentration (10 ng/ml) led to more profound changes compared to treatment with lower concentration (1 ng/ml). Taken together, these data support that changes in the chemo-physical environment associated with tissue injury or degeneration collectively impact chromatin organization in tenocytes and may instigate the phenotypic changes characteristic of loss of resident cell function.

### Degeneration and Aging Impacts Mechanical Sensitivity of Human Tenocytes

Given that human tenocytes rapidly respond to changes in their mechanical environments with alterations in chromatin spatial organization and condensation, we finally examined how tissue degeneration or aging impacts their mechanical sensitivity. To do this, young, tendinosis, or aged tenocytes were seeded onto the “stiffening” gel system and pre-cultured for 24 hours followed by additional 24 hours of culture after stiffening. Samples were then fixed to observe nanos-cale chromatin organization (**Fig. 6a**). Consistently, as was seen in soft substrates, H2B in young, tendinosis, and aged tenocytes were primarily localized to the nuclear periphery within 4 hours after stiffening (**Fig. 6b**). With 24 hours of stiffening, in young tenocytes (Young), chromatin became more uniformly dispersed and de-compacted throughout the nucleus (**Fig 6b-d**). Aged cells (Age) responded with the same trend to the increase in substrate stiffness, but to a lesser extent in terms of chromatin de-condensation (**Fig. 6b-d**). However, when the tendinopathic cells (Tendinosis) were cultured on the stiffening gels, no change in chromatin spatial organization was observed (**Fig. 6b-d**) and the extent of chromatin de-compaction, particularly at the nuclear periphery, was substantially lower compared to young and aged cells (**Fig. 6b-d**). These data suggest that the prolonged alterations in nano-scale chromatin organization in tenocytes may be associated with a loss of mechanical sensitivity with degeneration.

**Figure 6.**
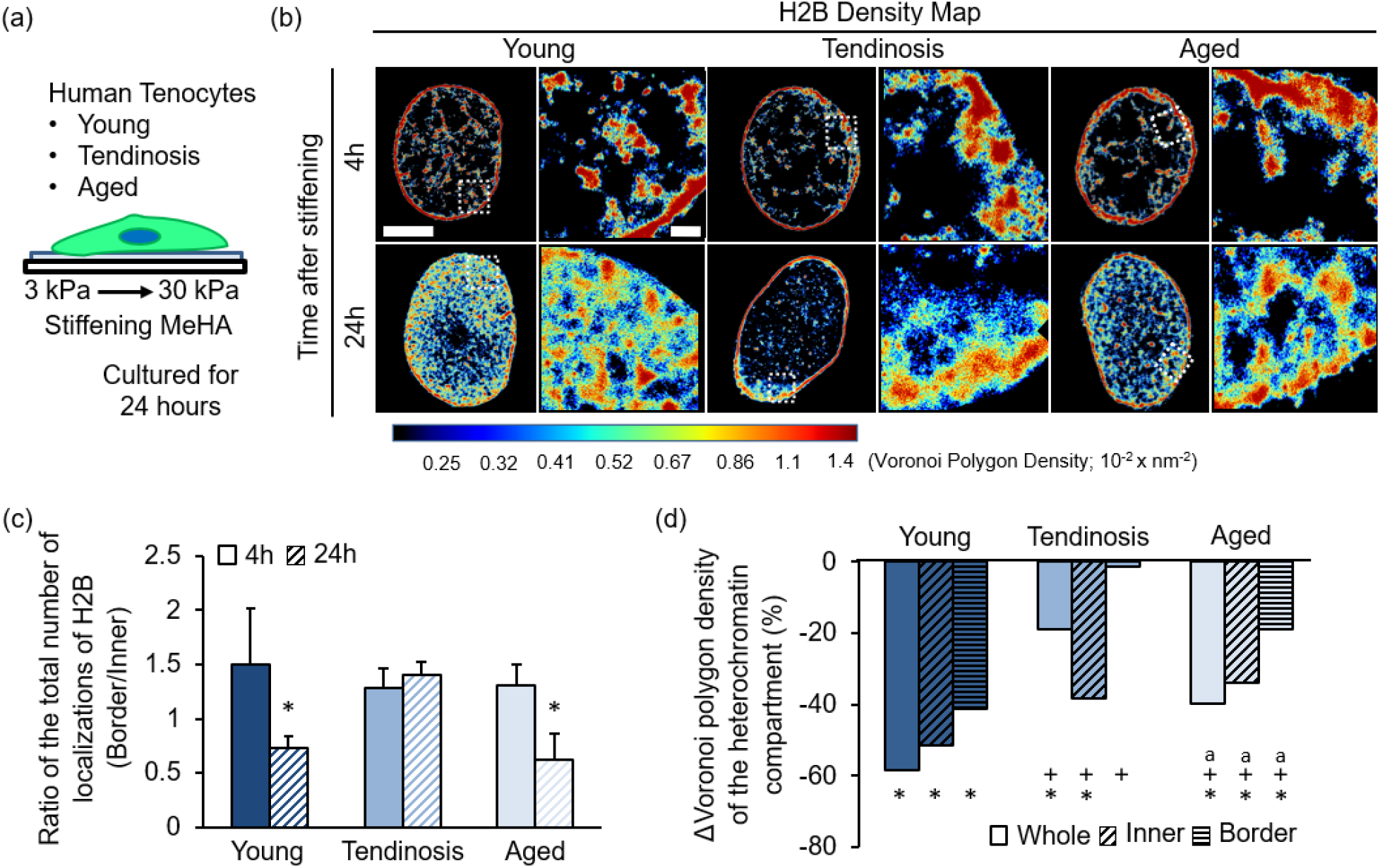
Altered mechanical sensitivity of human tenocytes with tissue degeneration and aging. (a) Schematic showing human young, tendinosis, or aged tenocytes seeded onto the “stiffening” hydrogel system and pre-cultured for 24 hours followed by an additional 24 hours of culture after stiffening. (b) Representative STORM super resolution images of H2B rendered as a density map showing changes to H2B localization in human young, tendinosis, or aged tenocytes 4 hours or 24 hours after stiffening. (c) Quantification of the ratio of the total number of H2B localizations per unit area at the nuclear border to the total number of H2B localizations in the inner region of the nucleus, showing a lack of response in the relocalization of H2B to the nuclear interior after stiffening in tendinosis tenocytes, and (d) changes in chromatin condensation in the condensed heterochromatin compartment 24 hours after stiffening. The Voronoi polygon density of the condensed heterochromatin compartment is shown normalized to the Voronoi polygon density 4 hours after stiffening (n = 5 nuclei/group, *: p<0.05 vs. 4h, +: p<0.05 vs. Young, a: p<0.05 vs. Tendinosis).

## Discussion

In this work, we demonstrated that nano-scale organization of chromatin within the nucleus of mesenchymal progenitor and adult fibrous tissue cells is highly responsive to mechanical perturbations, and that this responsivity depends on a patent cytoskeletal network and dynamic changes in histone modifications as well as the health/disease status of the cells themselves.

Using super-resolution microscopy, we first quantified how biophysical cues regulate nano-scale spatial chromatin organization in human mesenchymal progenitors. Our findings showed that substrate stiffness regulated both the spatial organization and the condensation state of chromatin at the nano-scale level, with stiff substrates resulting in less condensed chromatin and increased H3K4me3 nanodomains. Notably, these nanodomains were distributed throughout the MSC nuclei when compared to their localization on soft substrates. Given that such epigenetic marks and physical distribution is associated with active chromatin, this data suggested that stiff substrates may enhance the overall transcriptional activation of these cells. Conversely, and perhaps more surprisingly, when these same cells were placed on soft substrates, highly condensed chromatin domains relocalized to the nuclear periphery, a region in which constitutive heterochromatin resides and methyltransferases are abundant ^16,29^. This nuclear periphery has long been considered as a transcriptionally repressive area^40^ where inner nuclear membrane proteins in the nuclear lamina interact with proximal chromatin creating heterochromatin structures^41,42^. It has recently been suggested that this laminal sequestration at the nuclear envelope depends on specific epigenetic modifications, and indeed we observed that human MSCs on soft substrates increased their nano-scale H3K27me3 localization at the nuclear periphery. Notably, when the activity of EZH2 (a selective inhibitor of histone H3K27 methyltransferase) was inhibited, these spatial and epigenetic changes were prevented, even on soft substrates. These data indicate that soft substrates cause chromatin condensation and chromatin localization primarily to the nuclear periphery mediated by EZH2 likely resulting in genome-wide transcriptional suppression.

Having established that cells respond to alterations in their microenvironments by rapidly remodeling their genome organization, and further given that the development or degeneration of tissues alters biophysical properties of connective tissues^3–5^, we next investigated how tissue development or degeneration impact chromatin organization in tissue resident cells (i.e., tenocytes from tendon). Interestingly, our data show that tissue aging (which increases tissue stiffness^43,44^) decreased chromatin condensation in human tenocytes, while degeneration (which decreases local tissue stiffness^12,45^) increased chromatin condensation and lead to the relocalization of condensed chromatin to the nuclear periphery. Further, surprisingly, when healthy young tenocytes were exposed to degenerative chemo-physical environments, including low substrate stiffness^12,45^, low oxygen tension (which is altered with tissue injury and is critical in early tendinopathy^10,36,37^), and inflammation^38,39^, this bias in chromatin localization to the nuclear periphery was increased, similar to how chromatin is organized within the nucleus of degenerative tenocytes from tendinosis patients. Aberrant chromatin reorganization is associated with many diseases which leads to defects in gene regulation resulting in pathological gene expression programs^46–49^ and abnormal protein/matrix production of the cells. This work suggests that altered cellular microenvironments during development or degeneration, perhaps as a result of microscale damage and altered local chemo-physical environments, could lead to aberrant genome organization and differentiation of endogenous progenitor cells residing in the tissues. Most notably, when these degenerative cells were exposed to dynamic mechanical environments that restore physiologic chromatin architecture, they remained unresponsive. This suggests that when cells reside within a diseased state for a long enough period of time, the chromatin nano-material may undergo a permanent change. Further, the lack of mechano-responsivity in these degenerative cells may indicate that even if surgical and/or rehabilitative approaches are applied, the cells within these tissues will fail to resume their normal functionality.

Although this study shows fibrous connective tissue degeneration or “degenerative” chemo-physical cues cause nano-scale chromatin reorganization, the central question that remains unanswered is what specific genetic loci move to the periphery and the consequences of these spatial reorganization events on the transcriptional activity of such loci during degeneration. Given the massive chromatin re-localization and condensation that we observed here in degenerative tenocytes, it is likely that the effects are genome-wide and not restricted to a few specific genomic regions. In addition, given that histone modifications govern chromatin organization which is mediated by specific enzyme activities (i.e., histone methyltransferases or histone acetyltransferases), small molecule epigenetic modifiers may be considered to restore healthy chromatin structure and gene expression in degenerative cells to improve connective repair. Future work in this area should employ such strategies, with a specific focus on targeting key loci that are critical for tissue function and regeneration,

Overall, these findings expand our knowledge of the link between biophysical and chemical microenvironment and genome organization, opening the door for the identification of epigenetic factors that promote the recovery of a healthy chromatin structure in degenerative tenocytes to improve native fibrous connective tissue regeneration. This may also advance diagnostic and therapeutic approaches and novel regenerative medicine strategies in dense connective tissue repair.

## Acknowledgements

The authors would like to thank Dr. Kwang Hoon Song for assistance with the design of the microfluidic chamber, Dr. Elena Sorokina for assistance with the preparation of reagents for STORM imaging, Dr. Peter K. Relich for help with data visualization and analysis tools. The research was funded by the National Institutes of Health (K01 AR07787), the Penn Center for Musculoskeletal Disorders (P30 AR069619), and the Department of Veterans Affairs (IK6 RX003416), and the NSF Science and Technology Center for Engineering Mechanobiology (CMMI-1548571).

## Author Contributions

SJH, ST, XC, CL, BX, RM, JAB, VS, RLM and ML designed the studies. SJH, ST, XC, CL, and BX performed the experiments. SJH, ST, XC, CL, BX, RM, JAB, VS, RLM and ML analyzed and interpreted the data. SJH, RLM and ML drafted the manuscript, and all authors edited the final submission.

## Materials and Methods

### Preparation of cells

Human MSCs were isolated from fresh bone marrow from human donors (Lonza, male 23 years, female 18 years, male 22 years) as in ^50,51^. Briefly, using Ficoll density gradient centrifugation (800 RCF, 20 min), bone marrow was separated to obtain mononuclear cells, which were then plated and expanded on tissue culture plastic in alpha-modified essential medium (α-MEM, 10% FBS, 1% penicillin/streptomycin, 5 ng/ml basic fibroblast growth factor) at 37 °C/5% CO2 until 80% confluency of the colonies and stored in liquid nitrogen (95% FBS, 5% dimethylsulfoxide). All hMSCs were expanded in standard growth media (α-MEM, 10% FBS, 1% penicillin/streptomycin).

Human tenocytes were isolated from human finger flexor tendon tissues according to established protocols^10,52^. Human tendon was obtained from young (42 years), aged (81 years), or diagnosed with tendinosis (35 years) patients undergoing revision amputation for traumatic hand injury using IRB-approved protocols (#13D.238). Briefly, human tendon tissues (2 × 2 mm pieces) were digested in collagenase solution and the digested tissues were centrifuged. The cell pellets were resuspended and plated in basal growth media [high glucose DMEM, 10% pen/strep, L-glutamine, 10% FBS]. Experiments were performed using tenocytes of early passage (less than or equal to passage 6).

### Methacrylated hyaluronic acid synthesis

Methacrylated hyaluronic acid (MeHA) was synthesized as described previously^14,53,54^. Briefly, sodium hyaluronate (Lifecore, ∼74 kDa) was dissolved in deionized water (2wt%) and 1M NaOH added to adjust the pH to 9.5, followed by addition of methacrylic anhydride (MA, 2.8 mL per gram of HA). The reaction was allowed to proceed for 4 h on ice while maintaining a pH of 9.5 followed by a second addition of MA (2.8 mL of MA per gram HA) for 4 h at a pH of 9.5. The polymer was purified via dialysis, lyophilized and modification of ∼100% confirmed using ^1^H NMR.

### Methacrylated hyaluronic acid film fabrication and stiffening

To obtain MeHA hydrogel films (6 wt%) with an elastic modulus of ∼3 kPa (soft), MeHA was prepared using Michael-type addition crosslinking with dithiothreitol (0.97 mM) in PBS containing 0.2 M triethanolamine at pH 9)^14^. Hydrogel precursor solution containing thiolated RGD peptide (1 mM) were pipetted onto coverslips functionalized with 3-(trimethoxysilyl)propyl methacrylate and incubated for 30 min at 37 °C to obtain a crosslinked ∼15 µm thick hydrogel film^14^. To generate ‘stiffening hydrogels’ with an elastic modulus of ∼30 kPa, ‘soft hydrogel’ films were incubated in 0.05% lithium acylphosphinate (LAP) in PBS for 30 min at 37 °C followed by exposure to blue light (400 - 500 nm, 10 mW cm^−2^) for 5 min^32,33^. ‘Stiff hydrogels’ (∼30 kPa) were fabricated by exposing hydrogel precursor solutions in PBS containing 1 mM RGD and 0.05% LAP to blue light (400 - 500 nm, 10 mW cm^−2^) for 3 min.

### Theoretical modeling

In our modeling approach, the volume fractions of euchromatin, heterochromatin, and nucleoplasm that depend on the location ***x*** and time *t*, are defined as *ϕ*_*e*_(***x**,t*), *ϕ*_*h*_(***x**,t*), and *ϕ*_*n*_(***x**,t*), respectively, such that *ϕ*_*e*_+*ϕ*_*h*_+*ϕ*_*n*_ = 1. Motivated by the experimental H2B density maps (**Fig 3e & f**), we considered chromatin as phase-separating into a 1) a euchromatin-rich phase and 2) a heterochromatin-rich phase within the nucleus. Each of these phases contains different amounts of nucleoplasm. We construct a free energy density^55–57^, *f* as a function of volume fractions *ϕ*_*h*_ and *ϕ*_*e*_,

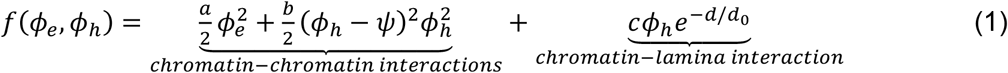

where the parameters *a* and *b* characterize the entropic contribution and enthalpy arising from chromatin-chromatin interactions. The double well defined by the second term captures the separation of chromatin into two phases, a heterochromatin-rich phase with *ϕ*_*h*_ = *ψ*, and a euchromatin-rich phase with *ϕ*_*h*_∼ 0. The last term captures interactions between heterochromatin and nuclear lamina, which gradually vanish over a length scale, *d*_0_,when the distance to the nuclear lamina, *d*, increases. The parameter *c* < 0 characterizes the strength of the chromatin-lamina interactions, with a preference for the heterochromatic phase at the lamina as suggested by previous studies^58–60^. The total free energy of the system can then be written as,

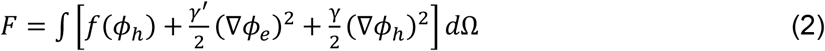

where the integral is over the volume of the nucleus, Ω. The terms proportional to *γ* and *γ*′ in the integrand penalize strong gradients of *ϕ*_h_ and *ϕ*_*e*_, which correspond to the energy of the interface between the heterochromatic and euchromatic phases (**SM Section 1**). The temporal evolution of the two phases of chromatin is governed by diffusion in response to the gradients in the chemical potential of eu- and hetero-chromatin 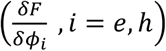, as well as the interconversion of the two phases due to methylation and acetylation regulated by epigenetic factors (**Fig 3a**). The evolution equations can be written as ^57,61,62^,

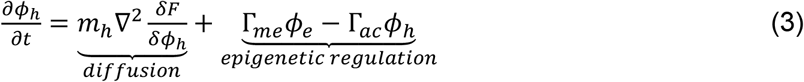

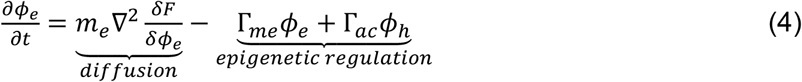

Here, *m*_*h*_ and *m*_*e*_ denote mobilities of heterochromatin and euchromatin. Note that while diffusion kinetics alone would individually conserve the total amount of heterochromatin and euchromatin in the nucleus, methylation and acetylation reactions allow conversion of euchromatin into heterochromatin and vice versa. For simplicity, we assume both acetylation and methylation follow first-order kinetics, although the terms can be made as complicated as needed. We use Γ*_me_* and Γ*_ac_* to denote the rates of methylation and acetylation, respectively. These two terms can capture the impact of the down (or up) regulation of enzymes such as histone deacetylases (HDACs) or histone acetyl transferases (HATs) that facilitate the switch in epigenetic states^63,64^. We set up the simulation to numerically solve Eq. 3-4 with parameters specified in **Supplementary Table S1**. We introduced small perturbations to *ϕ*_*h*_ and *ϕ*_*e*_ around their initial values to initiate the separation of chromatin into the two phases, and let the simulation proceed until a steady state was reached.

### Fabrication of microfluidic chamber

To apply fluid-induced shear stress, a microfluidic chamber system was developed. For this, a silicone elastomer base (Sylgard^™^ 184 PDMS) and curing agent were mixed thoroughly at a ratio of 10:1 (w/w) and the mixture was then degassed in a vacuum chamber at room temperature for 10 minutes. The cleared mixture was poured over 3D printed patterned acrylic masters (microfluidic channel dimensions: 5 mm (W) × 0.5 mm (H) × 20 mm (L) sitting on a petri dish. The device was cleared of bubbles and cured at 80°C for 1 h. PDMS molds were separated from the petri dish and the masters, and isolated from each other using a craft knife. The edges of the PDMS casts were chamfered to ensure exact flush between the cast surface and the coverslip glass. The inlet and outlet were drilled with a biopsy punch (D = 1 mm), concentric with the inlet and the outlet reservoirs. A thin layer of non-cured PDMS and curing agent mixture was applied around the microfluidic channel, and a coverslip glass was gently attached onto the bottom of the cast. The assembled chambers were left to cure at room temperature overnight before use.

### Stochastic optical reconstruction microscopy (STORM) imaging and analysis

For STORM imaging, cells were fixed in methanol-ethanol (1:1) at −20°C for 6 minutes followed by a 1-hour incubation in blocking buffer containing 10 wt% BSA (Sigma) in PBS as previously described^7^. Samples were incubated overnight with rabbit anti-H2B (1:50, Proteintech), rabbit anti-H3K4me3 (1:100, Thermo, MA5-11199), or rabbit anti-H3K27me3 (1:100, Thermo, PA5-31817) at 4°C. After repeated washing in PBS, samples were incubated with secondary antibodies custom labeled with activator-reporter dye pairs (Alexa Fluor 405-Alexa Fluor 647, Invitrogen) for STORM imaging as previously described^7^. All images were taken using a commercial STORM microscope system from ONI (Nanoimager S) fitted with a 100X, 1.4 NA oil immersion objective and sCMOS Hamamatsu Ocra FLash camera. A previously described imaging buffer was used to promote proper photoswitching of the AlexaFluor 647 (10 mM Cysteamine MEA [Sigma Aldrich, #30070-50G], Glox Solution: 0.5 mg/ml 1 glucose oxidase, 40 mg/ml 1 catalase [all Sigma], 10% Glucose in PBS)^65,66^. The 640 nm laser (1000 mW peak power) was used at 40% setting to excite the reporter dye (Alexa Fluor 647, Invitrogen) and to switch it to the dark state and the 405 nm laser was gradually increased to reactivate the Alexa Fluor 647 in an activator dye (Alexa Fluor 405)–facilitated manner and maintain a constant density of active fluorophores. Camera exposure time was 15 ms and 30,000 total frames were recorded per image. STORM image localizations were obtained using Nanoimager software (ONI) and rendered with custom-written software Insight3 (a kind gift of Bo Huang, UCSF, USA)^7^. For quantitative analysis, we used custom-written MATLAB codes. A previously described method was adapted that segments super-resolution images based on Voronoi tessellation of the fluorophore localizations^67,68^. Voronoi tessellation of a STORM image assigns a Voronoi polygon to each localization, such that the polygon area is inversely proportional to the local localization density^15^. The spatial distribution of localization of each nucleus is represented by a set of Voronoi polygons such that smaller polygon areas correspond to regions of higher density. The nuclear area was calculated by summing up all of the Voronoi polygon areas. The Voronoi polygons at the nuclear edge are extremely large and were omitted for all quantifications. Localization density (total number of localizations per nuclear area) of nuclei will vary from different mechanical/chemical treatments. To compare the chromatin organizational changes from different mechanical/chemical treatments, the localization density of the nuclei should be of similar range. To address this, the Voronoi polygon areas of each nucleus was divided by its mean Voronoi polygon area, such that the mean localization density of each nucleus is in reduced units of 1. The inverse of a reduced Voronoi polygon area associated with a localization, gives the reduced Voronoi density. All reduced Voronoi densities of each nucleus for each category of treatment were pulled together and plotted as cumulative distribution plots. A threshold was set such that the first 0∼40 percentile of the Voronoi density distribution was assigned to sparse euchromatin and the remaining 41∼80 percentile was assigned to condensed heterochromatin. Domains were segmented by grouping adjacent Voronoi polygons with areas less than a selected threshold and imposing a minimum of 3 localizations per domain criteria to generate the final segmented data set. The border and inside of the nucleus were cropped using a custom-made macro in ImageJ.

### Statistical analyses

Statistical analysis was performed using Student t tests or Analysis of variance (ANOVA) with Tukey’s honestly significantly different post hoc tests (GraphPad Prisms 8, La Jolla, CA). Results are expressed as the means ± SD. Differences were considered statistically significant at p < 0.05.

## Notes

### Competing Interest Statement

The authors have declared no competing interest.

