## Supplementary Information for "Chemo-Mechanical Cues Modulate Nano-Scale Chromatin Organization in Healthy and Diseased Connective Tissue Cells"

*: co-corresponding authors


### ***Supplementary Model Development and Supplemental Model (SM) Figures***

#### 1. Size of heterochromatin clusters increases with methylation levels ($\phi_n = 0, a = 0, k_e = 0, \psi = 1$)

We consider a single spherical droplet of heterochromatin of radius R in a relatively large region of euchromatin. To derive an analytical expression for the steady state radius and the density profile for hetero- and euchromatin, we discuss a simplified case where we ignore the nucleoplasm, such that $\phi_n = 0, a = 0, k_e = 0$, and $\phi_e + \phi_h = 1$, which corresponds to a two-phase system. While the nucleoplasm changes the trend quantitatively, the qualitative aspects of the scaling of the steady-state radius with the rates of methylation and acetylation continue to hold.

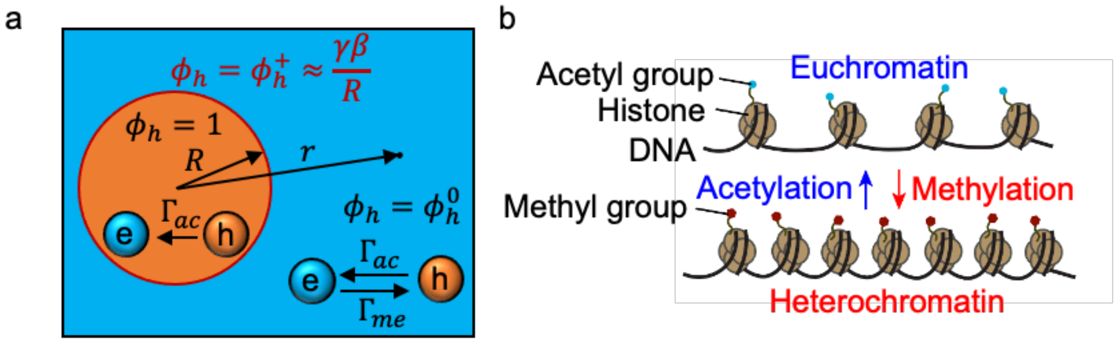


**Figure SM1** (a) Schematic of system of a single heterochromatin domain. (b) Conversion between heterochromatin and euchromatin through acetylation and methylation.

To analyze the steady-state size of a heterochromatin droplet, we first examine the concentration fields within and around the droplet. By taking advantage of spherical symmetry and using the polar coordinate with an origin at the center of the droplet (**Fig. SM1**), we can write the steady-state concentration field $\phi_h(r)$ as a function of distance to the center of the droplet, $r$. At the interface of the droplet (**Fig. SM1**), two different heterochromatin volume fractions $\phi_h^-$ and $\phi_h^+$ coexist inside and outside, respectively. The free energy associated with the interface yields the interfacial energy[1] $\kappa = \frac{1}{6}\sqrt{b\gamma}$, and

$$\phi_h^- \approx 1 \text{ and } \phi_h^+ \approx \frac{\kappa\beta}{R}, \tag{S1}$$

where the coefficient $\beta = \frac{2}{b}$ describes the effect of Laplace pressure on the volume fraction at the interface. We consider the case where the width of the interface between the two phases $w = 2\left(\frac{\gamma}{b}\right)^{1/2}$ is much smaller compared with the characteristic length for the variation of heterochromatin volume fraction $l = \left(\frac{m_h b}{\Gamma_{on}+\Gamma_{off}}\right)^{1/2}$. Outside the droplet, the phase is euchromatic with $\phi_h \ll 1$, and the free energy density function (Eq. 1) can be simplified to be $f(\phi_h) \approx \frac{b}{2}\phi_h^2$. Thus, under steady state conditions when the time derivatives vanish, Eq. 3 can be approximated as,

$$D_h \nabla^2 \phi_h - \Gamma_{ac}\phi_h + \Gamma_{me}\phi_e = 0. \tag{S2}$$

Here $D_h = m_h b$ is the diffusivity of heterochromatin. By solving Eq. S2 with boundary conditions $\phi_h|_{r=R} = \phi_h^+ = \frac{\kappa\beta}{R}$ and $\phi_h|_{r\to\infty} = \phi_h^0$ we get,

$$\phi_h = \phi_h^0 + \left(\phi_h^+ - \phi_h^0\right)\frac{R}{r}e^{(R-r)/l}, \tag{S3}$$

where $l = \left(\frac{m_h b}{\Gamma_{ac}+\Gamma_{me}}\right)^{1/2}$ is the characteristic reaction-diffusion length scale and $\phi_h^0$ is the volume fraction of heterochromatin far from the droplet. Because the droplet is small compared to the system, $\phi_h^0$ is dominated by the balance between the chemical reactions, such that,

$$\frac{d\phi_h^0}{dt} \approx \Gamma_{me}\left(1 - \phi_h^0\right) - \Gamma_{ac}\phi_h^0. \tag{S4}$$

At the steady state where $\frac{d\phi_h^0}{dt} = 0$, Eq. S4 yields $\phi_h^0 = \frac{\Gamma_{me}}{\Gamma_{me}+\Gamma_{ac}}$.

Having obtained the expression for the concentration field $\phi_h(r)$, we next discuss the dynamics of the growth of the droplet. The volume of the heterochromatic droplet $V$ can change due to 1) diffusive flux of heterochromatin from the euchromatic region, and 2) acetylation that converts heterochromatin to euchromatin. Therefore,

$$\frac{dV}{dt} = 4\pi R^2\frac{dR}{dt} = \underbrace{J(R)}_{diffusive\ flux} - \underbrace{\frac{4}{3}\pi R^3\Gamma_{ac}}_{acetylation}. \tag{S5}$$

The flux through a spherical shell with radius R is given by,

$$J = -4\pi R^2 D_h \frac{\partial\phi_h}{\partial r}. \tag{S6}$$

The spatial gradient $\partial\phi_h/\partial r$ in Eq. S6 can be approximated using Eq. S3 for a small droplet in a large system in the limit $R \ll l$. In the vicinity of the droplet interface ($r \to R$), $e^{(R-r)/l} = 1$, and therefore $\frac{\partial\phi_h}{\partial r} \approx (\phi_h^0 - \frac{\kappa\beta}{R})/R$. Combining Eq. S3-S6, we obtain a simplified equation for the dynamics of growth of droplet,

$$\frac{dR}{dt} = D_h\left(\frac{\phi_h^0}{R} - \frac{\kappa\beta}{R^2}\right) - \frac{\Gamma_{ac}}{3}R. \quad \text{(S7)}$$

Setting $\frac{dR}{dt} = 0$ in Eq. S7 yields the critical and stable steady-state radius of droplet (**Fig. SM2**).

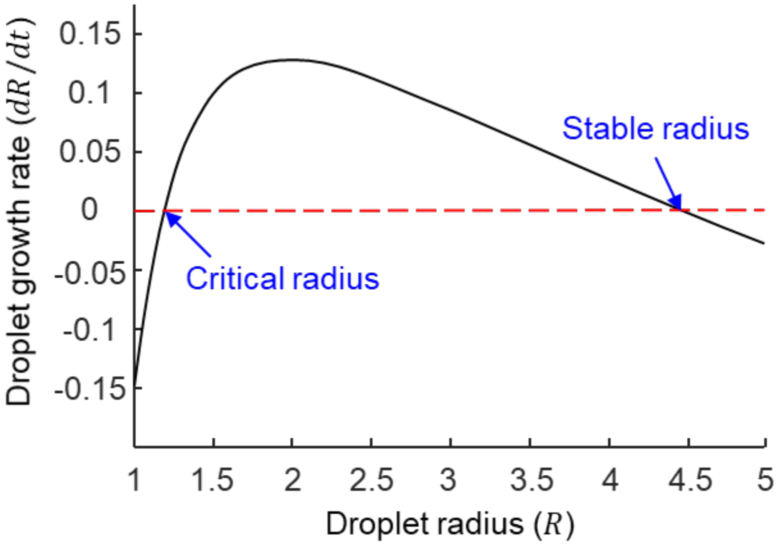


**Figure SM2** Growth rate of the droplet as a function of the droplet radius R in a large system. Above the critical radius, the heterochromatic droplet will grow in size. When the size of the droplet is larger than the stable radius, the droplet shrinks, and in steady state, the droplet remains at the stable radius. In the case of small interfacial energy $\kappa \ll R_S\phi_h^0/\beta$, the stable radius can be approximated by,

$$R_s = \sqrt{\frac{3D_h}{\Gamma_{me}+\Gamma_{ac}}\frac{\Gamma_{me}}{\Gamma_{ac}}}. \quad \text{(S8)}$$

Eq. S8 shows the scaling relation between the size of the heterochromatin and the methylation and acetylation levels.

This derivation can be easily repeated for the 2D case. Eq. S2 can be rewritten as $D_h\left(\frac{\partial^2\phi_h}{\partial r^2} + \frac{1}{r}\frac{\partial\phi_h}{\partial r}\right) - (\Gamma_{ac} + \Gamma_{me})\phi_h + \Gamma_{me} = 0$, which is the modified Bessel's equation. The solution to this equation is

$$\phi_h = \phi_h^0 + \left(\phi_h^+ - \phi_h^0\right)\frac{K_0(r/l)}{K_0(R/l)} \;, \quad \text{(S9)}$$

where $K_0$ denotes the Bessel K function. Also, in 2D Eq. S5 becomes

$$\frac{dV}{dt} = 2\pi R\frac{dR}{dt} = \underbrace{-2\pi R D_h\frac{\partial\phi_h}{\partial r}}_{diffusive\ flux} - \underbrace{\pi R^2\Gamma_{ac}}_{acetylation} \;. \quad \text{(S10)}$$

Solving Eq. S10 at the steady state $\frac{dV}{dt} = 0$ yields a scaling relation between the radius of heterochromatin domain and the level of methylation, $\frac{K_1(R_s/l)}{K_0(R_s/l)} \sim \frac{2D_h}{\Gamma_{me}+\Gamma_{ac}}\frac{\Gamma_{me}}{\Gamma_{ac}}$. Thus, we obtain the same scaling relation that $R_s = f\left(\frac{D_h}{\Gamma_{me}+\Gamma_{ac}}\frac{\Gamma_{me}}{\Gamma_{ac}}\right)$, where $f$ is a function determined from Eq. S10. In the range of our parameters, $R_s \sim \sqrt{\frac{D_h}{\Gamma_{me}+\Gamma_{ac}}\frac{\Gamma_{me}}{\Gamma_{ac}}}$ provides a good approximation for the scaling of the size of heterochromatin domains (**Fig 3c**).

### 2. Size of heterochromatin domain at the border of the nucleus increases with methylation levels

Heterochromatin can be physically tethered (for example through HDAC3) or chemically bound to the nuclear lamina. Such interactions between heterochromatin and the lamina[2] induce a preferential localization of heterochromatin domains near the border of the nucleus. To model this phenomenon, we explicitly introduce a heterochromatin-lamina interaction potential of the form $V = c\phi_h e^{-d/d_0}$ ($c < 0$) in the free energy density (Eq. 1), such that the energy of the system decreases as heterochromatin localizes to the border. The heterochromatin localization at the border of the nucleus is governed by the energetic competition between the chromatin-lamina interactions and chromatin-chromatin interactions, analogous to wall-wetting phenomenon described by the well-known Young's equation

$$\gamma_{le} = \gamma_{eh} cos\theta + \gamma_{lh}, \tag{S11}$$

where $\gamma_{le}$ denotes the lamina-euchromatin interfacial energy, $\gamma_{lh} = \gamma_{le} - V$ denotes the lamina-heterochromatin interfacial energy (increases as $c$ becomes more negative), and $\gamma_{eh}$ denotes the heterochromatin-euchromatin interfacial energy. $\theta$ denotes the contact angle. When the heterochromatin-lamina interaction is small (magnitude of $c$ is small), heterochromatin will form small discrete droplets at the wall such that contact angle $\theta$ is large (**Fig. S3**). Increasing the heterochromatin-lamina interfacial energy ( $\gamma_{lh}$ ) causes the contact angle between heterochromatin domains and lamina to decrease ($\theta \downarrow$), and their radius to increase ($R \uparrow$). When the heterochromatin-lamina interactions ( $c\phi_h e^{-d/d_0}$ ) are sufficiently strong ( $\theta \to 0, R \to \infty$ ), heterochromatin forms a continuous layer covering the lamina (**Fig. SM3 & SM4**). In our experiments, near the border of the nucleus, we observed a similar transition from sparsely distributed heterochromatin droplets to a continuous layer when cells were cultured on soft substrates (**Fig 3e & d**).

In addition to the heterochromatin-lamina interactions, the size of the heterochromatin domain at the boundary of the nucleus is also related to the acetylation and methylation levels. Here we derive the scaling relation between the thickness of heterochromatin domain and the methylation level.

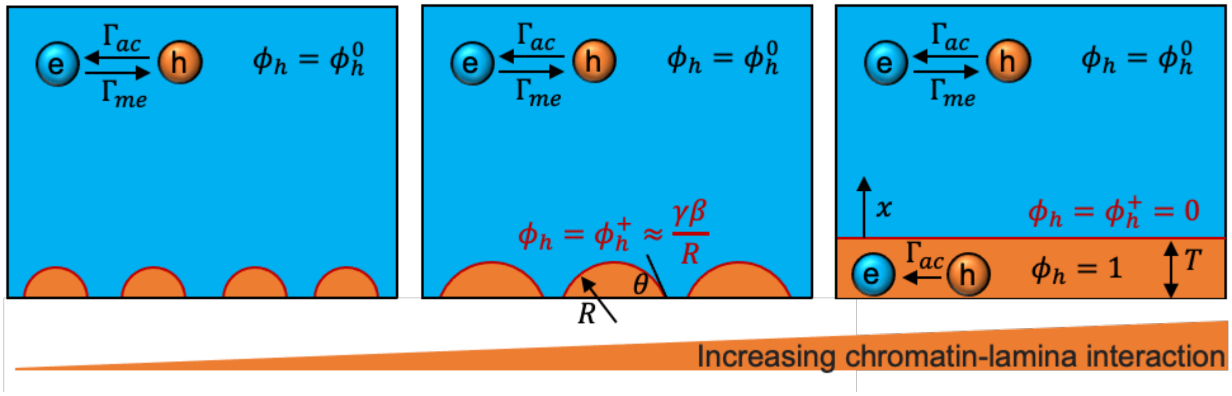


**Figure SM3** Schematics showing the formation of heterochromatin domains at the nucleus border.

The reaction-diffusion equation Eq. S2 can be simplified to an 1D equation,

$$D_h \frac{d^2\phi_h}{dx^2} - \Gamma_{ac}\phi_h + \Gamma_{me}\phi_e = 0, \tag{S12}$$

where $x$ denotes the distance from the interface between heterochromatin and euchromatin (**Fig. SM3**). Two different volume fractions $\phi_h^-$ and $\phi_h^+$ coexist immediately inside and outside the boundary of heterochromatin, respectively. Solving Eq S10 with boundary conditions $\phi_h|_{x=0} = \phi_h^+ = 0$ (let $R \to \infty$ in Eq. S1) and $\phi_h|_{x\to\infty} = \phi_h^0 = \frac{\Gamma_{me}}{\Gamma_{ac}+\Gamma_{me}}$ gives,

$$\phi_h = -\phi_h^0 e^{-x/l} + \phi_h^0 \quad \text{(S13)}$$

Similar to Eq. S5, the change in the volume of the layer of heterochromatin can be written as,

$$\frac{dV}{dt} = S\frac{dT}{dt} = SD_h\frac{\partial\phi_h}{\partial x} - ST\Gamma_{ac}, \quad \text{(S14)}$$

where S denotes the surface area of the heterochromatin layer. Solving Eq. S27 at the steady state by setting $\frac{dT}{dt} = 0$ yields

$$T = \frac{D_h\phi_h^0}{\Gamma_{ac}l} \sim \frac{\Gamma_{me}}{\Gamma_{ac}}\sqrt{\frac{1}{\Gamma_{me}+\Gamma_{ac}}}. \quad \text{(S15)}$$

The relation in Eq. is found to be true for the 2D case as well (S in Eq. S14 denotes a line instead of a surface). In **Fig. 3d** in the main text, we show excellent agreement between our analytically derived scaling relation and numerical simulations.

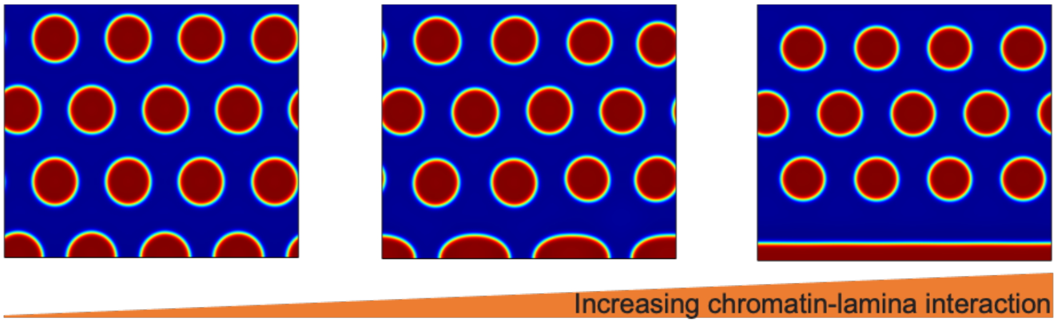


**Figure SM4** Transition of morphology of heterochromatin domains from discrete islands to continuous layer at the nucleus border due to increasing chromatin-lamina interaction.

#### 3. Table of parameters

The parameters are estimated such that the simulation can generate similar patterns of heterochromatin domains observed in our experiments. In this study, we focus on predicting general trends, and our quantitative analysis are presented with scaling relations rather than exact numbers. Only the methylation and acetylation levels $\Gamma_{me}$ and $\Gamma_{ac}$ are tuned to fit the experimental observations.

**Table SM1** Model parameters used in this study.

| Parameter | Value | Reference |
|---|---|---|
| Initial $\phi_h$ | 0.5 | Estimated from experimental H2B density |
| Initial $\phi_e$ | 0.5 | Estimated from experimental H2B density |
| Initial $\phi_n$ | $1 - \phi_e - \phi_h$ | By definition |
| $a$ | 0 | Estimated to initiate phase separation |

| | | |
|---|---|---|
| $b$ | 1 | Estimated to initiate phase separation |
| $c$ | 0.02 | Estimated |
| $d_0$ | 0.1 | Estimated |
| $\gamma'$ | 0 | Estimated |
| $\gamma$ | 1 | Estimated |
| $m_e$ | 0.04 | Estimated |
| $m_h$ | 0.04 | Estimated |
| $\Gamma_{me}$ | 0.21 (stiff) 0.2 (soft) 0.1 (GSK) | Fit to experiment |
| $\Gamma_{ac}$ | 0.7 (stiff) 0.4 (soft) 0.4 (GSK) | Fit to experiment |

### ***Supplementary Figures***

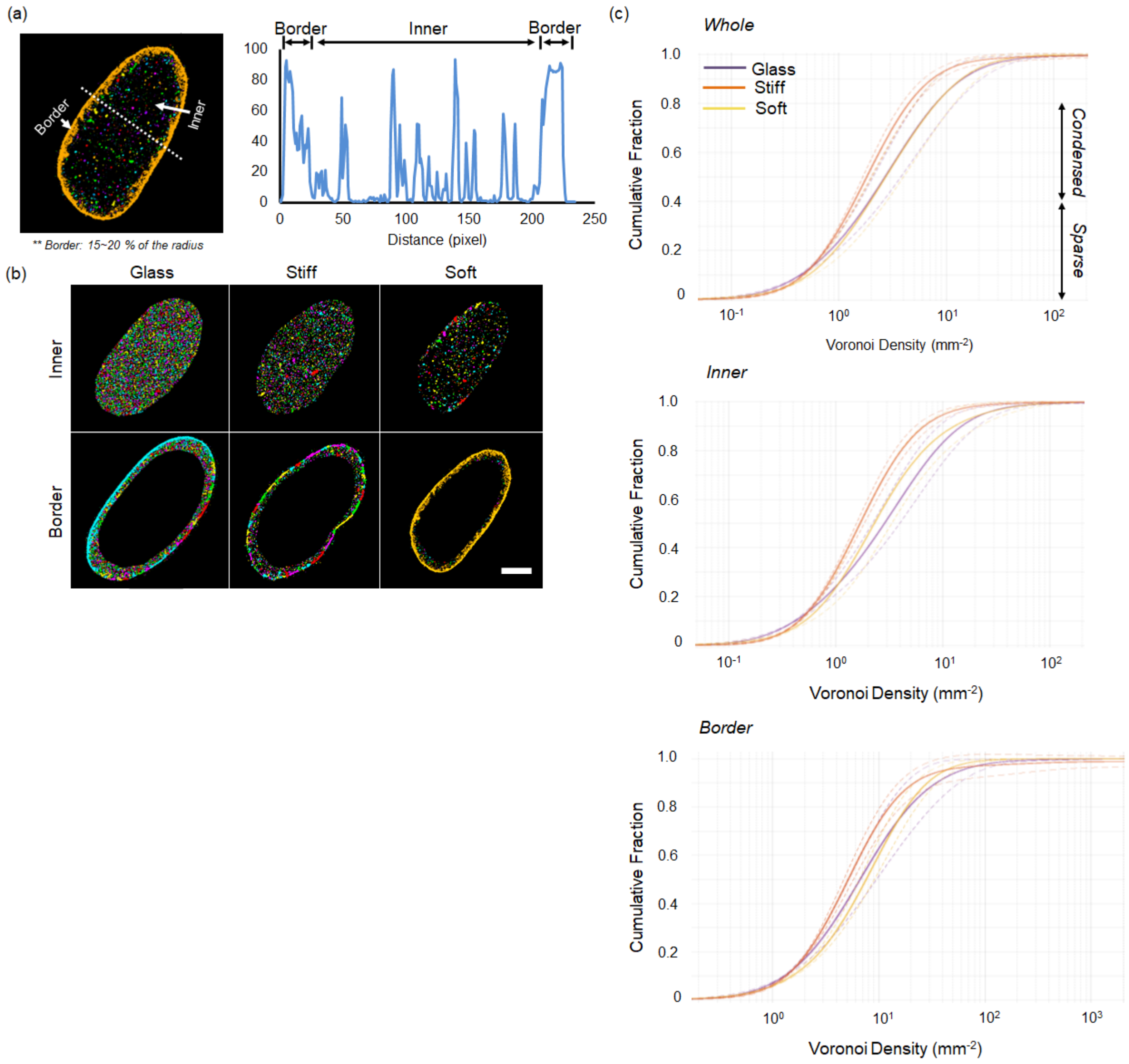


**sFigure 1.** (a) The nuclear periphery (Border) is defined as 15 ~ 20% of the outer area from the nuclear border determined from the STORM super-resolution image intensity profile across the nucleus. (b) Representative images showing pseudo-color-coded H2B nano-domains at nuclear border or inner part of the nucleus after splitting the nucleus into Border and Inner based on the analysis described in (a). (c) Cumulative distribution of Voronoi polygon density (inverse of Voronoi polygon area) averaged over all nuclei (solid lines) and shown for the whole, border part or inner part of hMSC nuclei on different substrates (dashed lines: standard deviation). Chromatin was split into two compartments, sparse euchromatin having low Voronoi density (0-40th percentile of the cumulative Voronoi polygon density plot) and condensed heterochromatin having high Voronoi polygon density (41-80th percentile of the cumulative Voronoi polygon density).

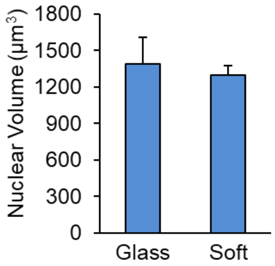


**sFigure 2.** Nuclear volume of hMSCs cultured on glass or soft substrate (n = 17 ~ 24).

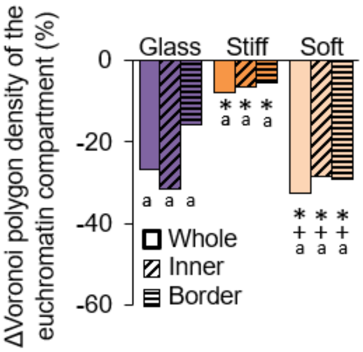


**sFigure 3.** Changes in chromatin condensation with the GSK treatment in sparse euchromatin compartment in hMSCs. The Voronoi polygon density of the sparse euchromatin compartment in GSK treated cells is shown normalized to control cells, showing a decrease in chromatin condensation of euchromatin with GSK treatment (n = 5 nuclei/group, a: $p<0.05$ vs. Ctrl, *: $p<0.05$ vs. Glass, +: $p<0.05$ vs. Stiff).

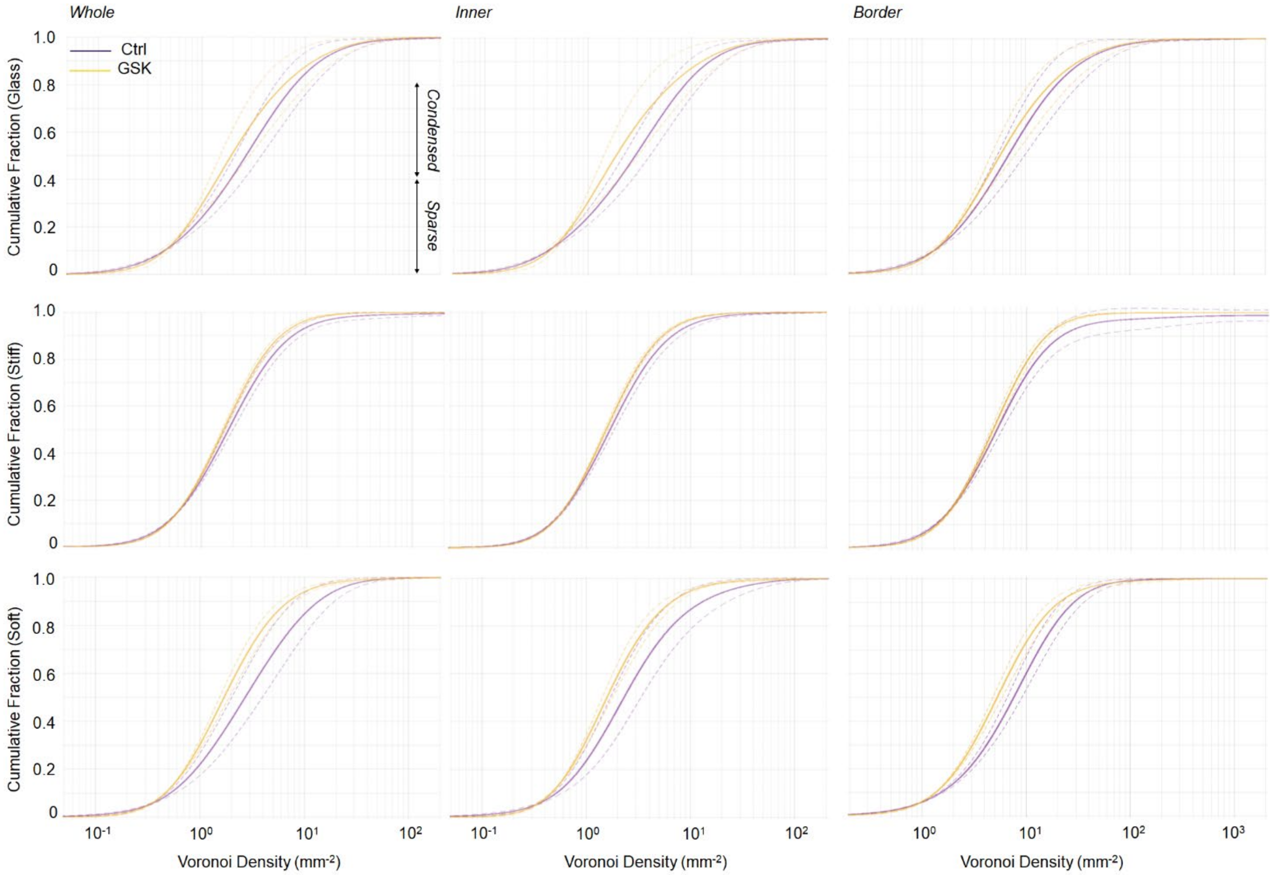


**sFigure 4.** Cumulative distribution of Voronoi polygon density (inverse of Voronoi polygon area) averaged over all nuclei and shown for the whole, the border, or inner portion of hMSCs nucleus on different substrates with/without GSK343 treatment (Ctrl or GSK). Solid lines: average and dashed line: standard deviation.

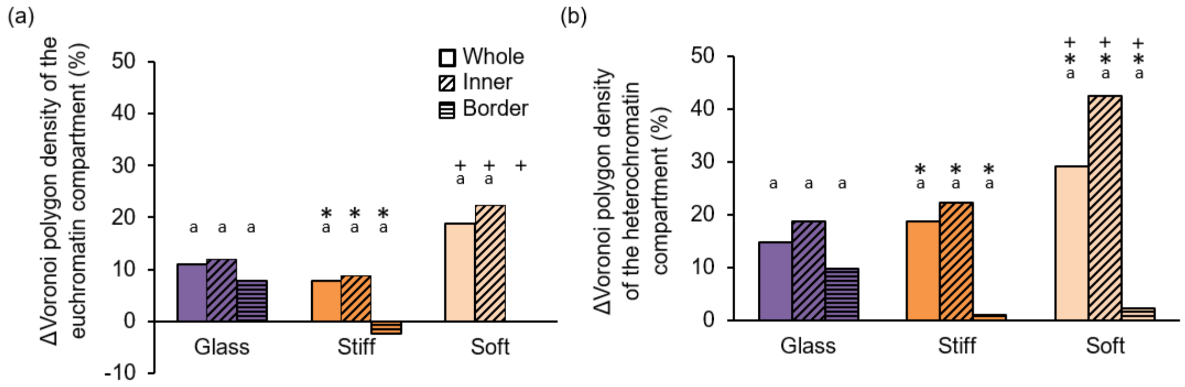


**sFigure 5.** Changes in chromatin condensation with the LPA treatment in sparse euchromatin or condensed heterochromatin compartments calculated for the whole, the border part, or inner of the nucleus in hMSCs cultured on glass, stiff or soft hydrogels. The Voronoi polygon density of the condensed heterochromatin or sparse euchromatin compartment in LPA treated cells is shown normalized to control cells, showing an increase in chromatin condensation of heterochromatin and euchromatin with LPA treatment (n = 5 nuclei/group, a: $p<0.05$ vs. Ctrl, *: $p<0.05$ vs. Glass, +: $p<0.05$ vs. Stiff).

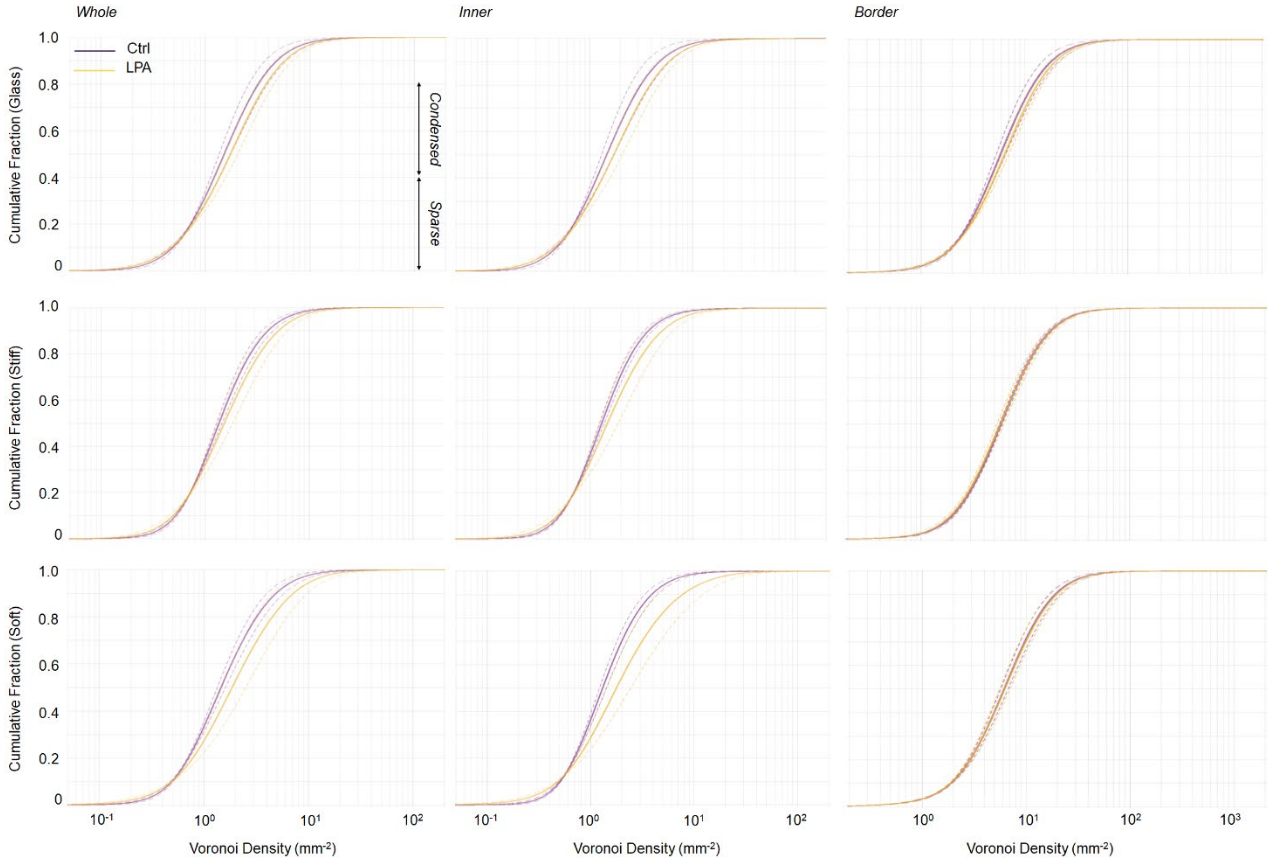


**sFigure 6.** Cumulative distribution of Voronoi polygon density (inverse of Voronoi polygon area) averaged over all nuclei and shown for the whole, the border, or inner region of hMSCs nucleus on different substrates with/without LPA treatment (Ctrl or LPA). Solid lines: average and dashed line: standard deviation.

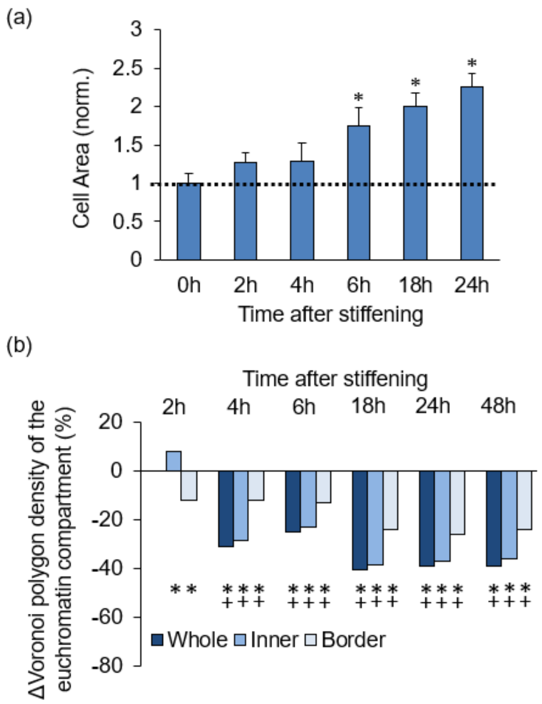


**sFigure 7.** (a) Changes in cell area in response to substrate stiffening between 0-24 hours after stiffening. Values are normalized to the starting cell area at 0h (n = 5 cells/group, normalized to the cell area at 0h, *:$p<0.05$ vs. 0h). (b) Changes in chromatin condensation in the sparse euchromatin compartment in hMSCs between 0-48 hours after stiffening. The Voronoi polygon density of the sparse euchromatin compartment in cells grown on the stiffening hydrogel is shown normalized to the starting Voronoi polygon density at 0h, showing a decrease in chromatin condensation of euchromatin with time after stiffening (n = 5 nuclei/group, *: $p<0.05$ vs. 0h, +: $p<0.05$ vs. 2h).

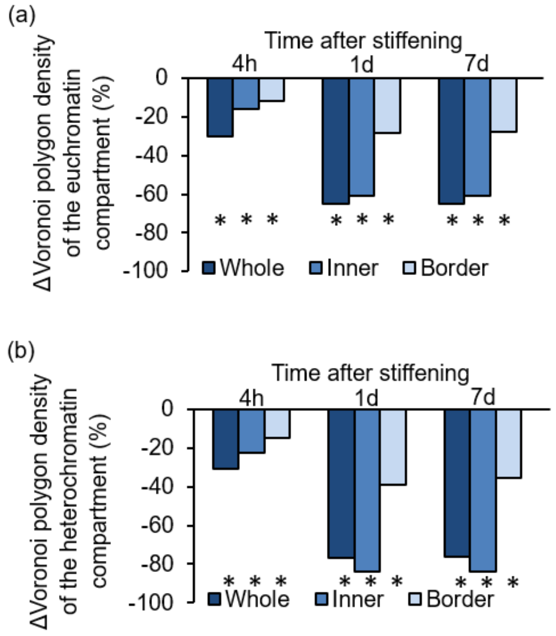


**sFigure 8.** Changes in chromatin condensation in sparse euchromatin (a) or condensed heterochromatin (b) compartments in hMSCs between 0-7 days after stiffening. The Voronoi polygon density of the condensed heterochromatin or the sparse euchromatin compartment in cells grown on the stiffening hydrogel is shown normalized to the starting Voronoi polygon density at 0h, showing a decrease in chromatin condensation with time after stiffening (n = 5 nuclei/group, *: $p<0.05$ vs. 0h, +: $p<0.05$ vs. 1d).

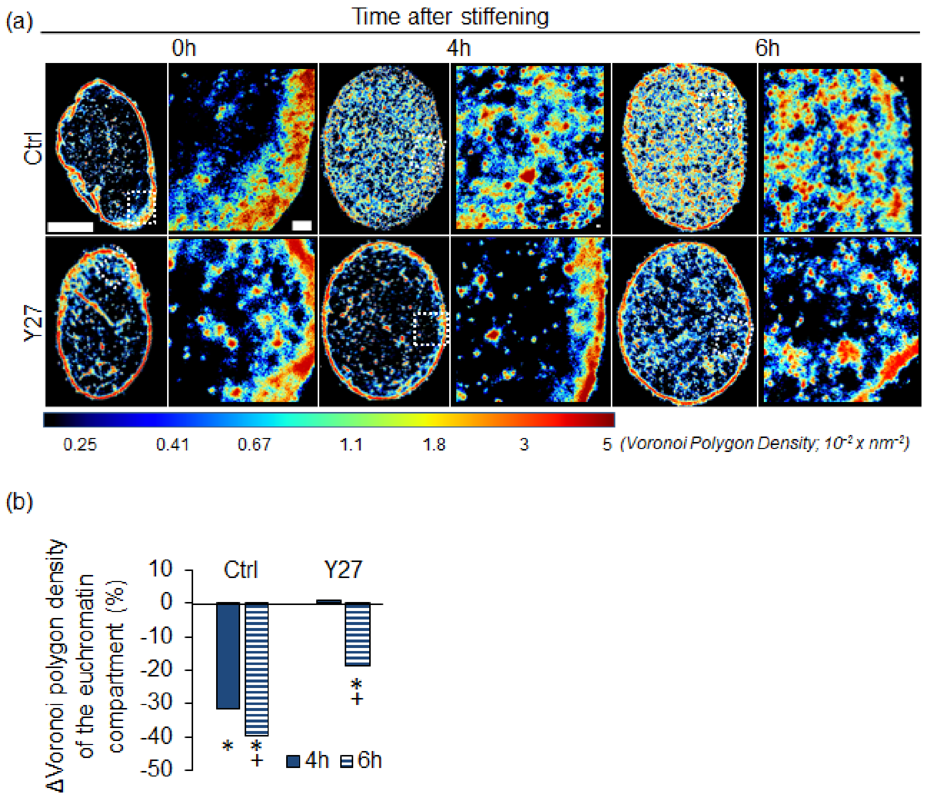


**sFigure 9.** (a) Representative STORM super-resolution image of H2B rendered as a density map showing chromatin redistribution between 0-6 hours after substrate stiffening [bars = 5um (left), 300 nm (right)] and qualifications of (b) changes in chromatin condensation in sparse euchromatin compartments in hMSCs between 0-6 hours after stiffening with/without the Y27 treatment. The Voronoi polygon density of the sparse euchromatin compartment in cells grown on the stiffening hydrogel with or without Y27 treatment is shown normalized to the starting Voronoi polygon density at 0h, showing that Y27 treatment prevents the decrease in chromatin condensation of euchromatin with time after stiffening (n = 5 nuclei/group, *: $p<0.05$ vs. 0h, +: $p<0.05$ vs. 4h).

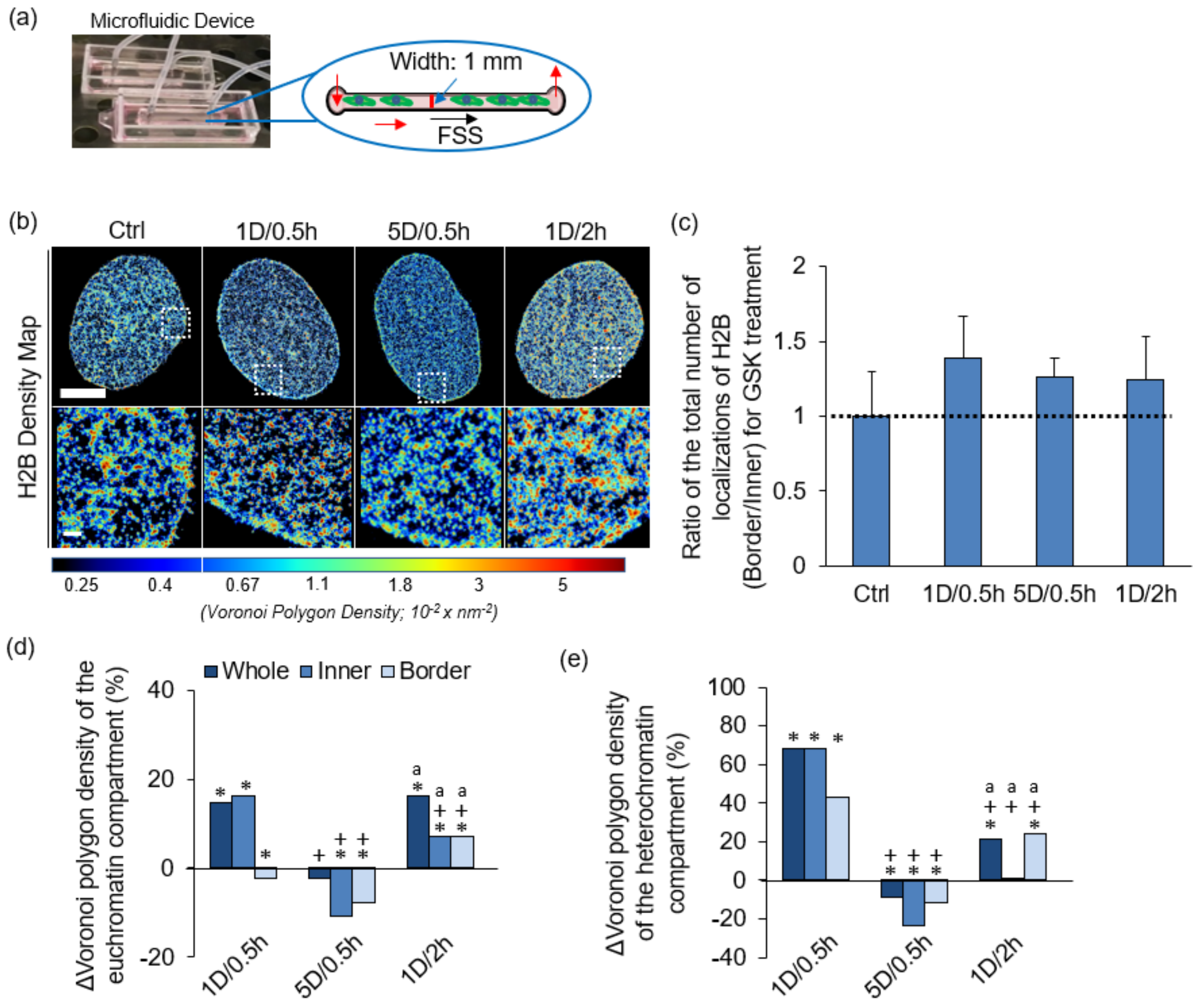


**sFigure 10.** (a) Picture and schematic of a custom-PDMS microfluidic fluid shear stress (FSS) device. (b) Representative STORM super-resolution images of H2B rendered as a density map showing redistribution of H2B in hMSCs in response to FSS of varying magnitude 1 ~ 5 dyne/cm$^2$) and duration (0.5 ~ 2 hours). (c) Quantification of changes in the ratio of the total number of H2B localizations per unit area at the nuclear border to the total number of H2B localizations at the inner part of the nucleus with/without the application of FSS (n = 5 nuclei/group, *: p<0.05 vs. Ctrl, +: p<0.05 vs. 1D/0.5h, a: p<0.05 vs. 5D/0.5h). (d) Changes in chromatin condensation with the application of FSS in (d) sparse euchromatin compartments or (e) condensed heterochromatin compartments in hMSCs. The Voronoi polygon density of the condensed heterochromatin or the sparse euchromatin compartment in cells subjected to FSS is shown normalized to the Voronoi polygon density in the absence of FSS (n = 5 nuclei/group, *: p<0.05 vs. Ctrl, +: p<0.05 vs. 1D/0.5h, a: p<0.05 vs. 5D/0.5h).

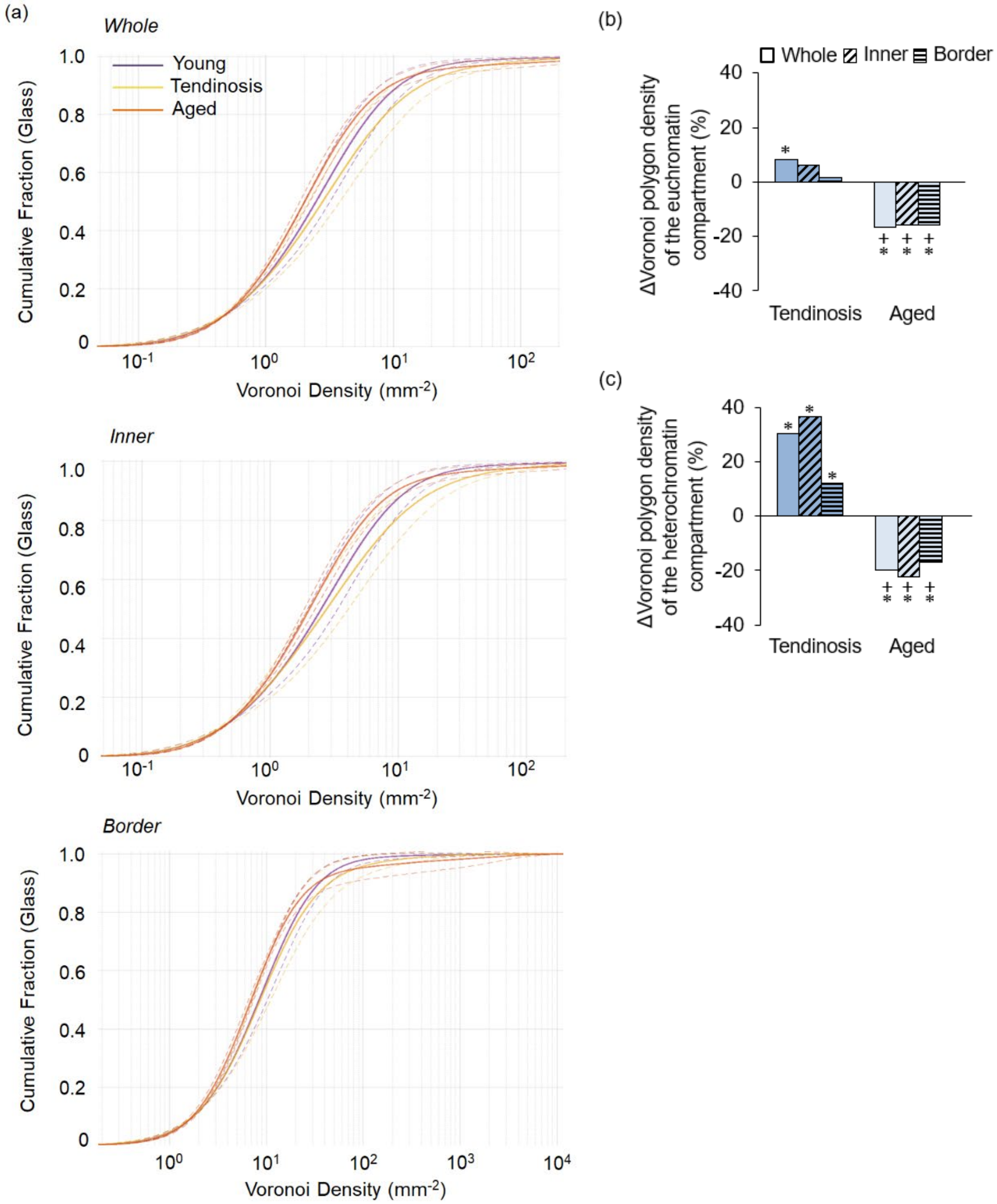


**sFigure 11.** (a) Cumulative distribution of Voronoi polygon density (inverse of Voronoi polygon area) in the whole, the border, or inner region of the nucleus averaged over all nuclei of Young, Tendinosis, or Aged human tenocytes cultured on cover glasses (solid lines: average, dashed lines: standard deviation, n = 5 nuclei/group, *: p<0.05 vs. Young, +: p<0.05 vs. Tendinosis). Changes in chromatin condensation in the sparse euchromatin compartment (b) or condensed heterochromatin (c) of human tenocytes from young and tendinosis and aged donors. The Voronoi polygon density of the euchromatin or heterochromatin compartment is shown normalized to the Voronoi polygon density in human tenocytes obtained from young and healthy donor (n = 5 nuclei/group, *: p<0.05 vs. Young, +: p<0.05 vs. Tendinosis).

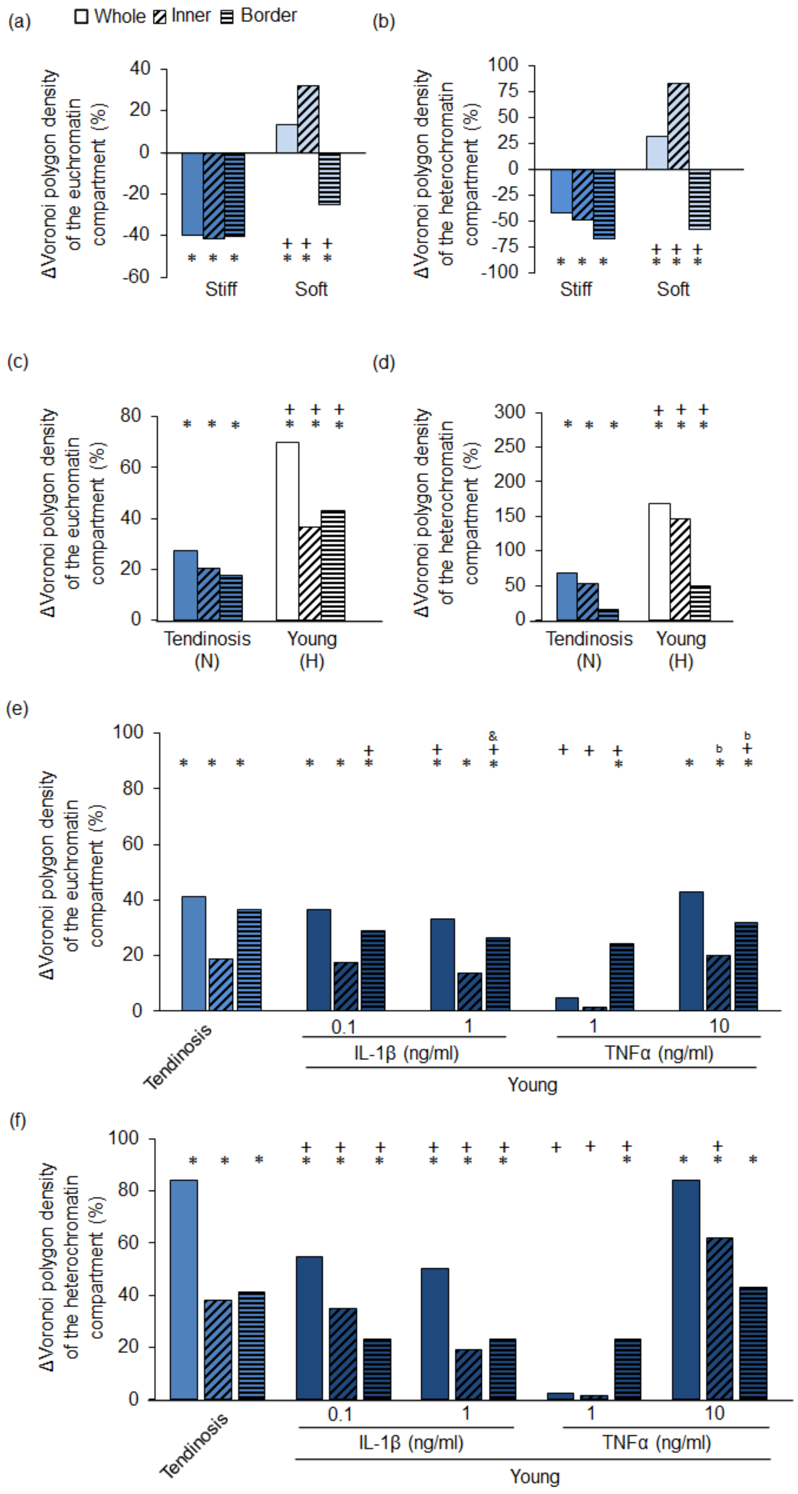
(a)
Whole
Inner
Border
ΔVoronoi polygon density of the euchromatin compartment (%)
Stiff
Soft
(b)
ΔVoronoi polygon density of the heterochromatin compartment (%)
Stiff
Soft
(c)
ΔVoronoi polygon density of the euchromatin compartment (%)
Tendinosis (N)
Young (H)
(d)
ΔVoronoi polygon density of the heterochromatin compartment (%)
Tendinosis (N)
Young (H)
(e)
ΔVoronoi polygon density of the euchromatin compartment (%)
Tendinosis
IL-1β (ng/ml)
TNFα (ng/ml)
Young
(f)
ΔVoronoi polygon density of the heterochromatin compartment (%)
Tendinosis
IL-1β (ng/ml)
TNFα (ng/ml)
Young

**sFigure 12.** Changes in chromatin condensation with substrate stiffness in sparse euchromatin (a) or condensed heterochromatin (b) compartments. The Voronoi polygon density of the sparse or condensed heterochromatin compartment is shown normalized to the Voronoi polygon density in human tenocytes grown on Glass, showing a decrease/increase in the condensation of heterochromatin compartment on stiff/soft substrates (n = 5 nuclei/group, *: $p<0.05$ vs. glass, +: $p<0.05$ vs. soft). Changes in chromatin condensation in sparse euchromatin (c) or condensed heterochromatin (d) compartments. The Voronoi polygon density of the sparse or condensed heterochromatin compartment is shown normalized to the Voronoi polygon density in human young tenocytes grown under normoxic (N) conditions, showing an increase in the condensation of heterochromatin under hypoxic (H) conditions (n = 5 nuclei/group, *: $p<0.05$ vs. young healthy tenocyte under the normoxic conditions, +: $p<0.05$ vs. tendinosis tenocyte under the normoxic conditions). Changes in chromatin condensation with exposure to inflammatory cytokines in sparse euchromatin (e) or condensed heterochromatin (f) compartments. The Voronoi polygon density of the sparse or condensed heterochromatin compartment is shown normalized to the Voronoi polygon density in human young tenocytes without treatment with inflammatory cytokines, showing an increase in the condensation of heterochromatin upon treatment (n = 5 nuclei/group, *: $p<0.05$ vs. young human tenocyte control, +: $p<0.05$ vs. tendinosis young human tenocyte).
